# Loss of the lysosomal lipid flippase ATP10B leads to progressive dopaminergic neurodegeneration and Parkinsonian motor deficits

**DOI:** 10.1101/2024.07.09.602700

**Authors:** María Sanchiz-Calvo, Elena Coccia, Christopher Cawthorne, Gustavo Morrone Parfitt, Koen Van Laere, Teresa Torre-Muruzabal, Diego Cabezudo, George Tsafaras, Ana Cascalho, Chris Van den Haute, Peter Vangheluwe, Joel Blanchard, Eduard Bentea, Veerle Baekelandt

**Author notes:** Corresponding Author Baekelandt, Veerle.

## Abstract

**Background:** ATP10B, a transmembrane lipid flippase located in late endosomes and lysosomes, facilitates the export of glucosylceramide and phosphatidylcholine by coupling this process to ATP hydrolysis. Recently, loss-of-function mutations in the *ATP10B* gene have been identified in Parkinson’s disease patients, pointing to *ATP10B* as a candidate genetic risk factor. Previous studies have shown compromised lysosomal functionality upon *ATP10B* knockdown in human cell lines and primary cortical neurons. However, its role *in vivo* and specifically in the nigrostriatal dopaminergic system remains poorly understood.

**Methods:** To investigate the role ATP10B in PD neuropathology, we induced ATP10B knockdown specifically in *substantia nigra pars compacta* neurons of rats using viral vector technology. Two different microRNA-based shRNA constructs targeting distinct regions of the ATP10B mRNA were used to cross-validate the findings. Behavioral evaluation, dopamine transporter ^18^F-FE-PE2I positron emission tomography imaging and neuropathological examination of the nigrostriatal pathway at one year post-injection were conducted. Additionally, midbrain neuronal cultures derived from ATP10B knock-out human induced pluripotent stem cells clones were used to study the impact of ATP10B loss in dopaminergic neurons in a more translational model.

**Results:** *ATP10B* knockdown in rat brain induced Parkinsonian motor deficits, and longitudinal striatal dopamine transporter ^18^F-FE-PE2I PET imaging revealed a progressive decrease in binding potential. Immunohistochemical analysis conducted one year post-injection confirmed the loss of dopaminergic terminals in the *striatum*, alongside a loss of dopaminergic neurons in the *substantia nigra pars compacta*. The expression of LAMP1, LAMP2a, cathepsin B and glucocerebrosidase was studied by immunofluorescence in the surviving dopaminergic neurons. A decrease in lysosomal numbers and an increase in lysosomal volume were observed more consistently in one of the knockdown constructs. The vulnerability of dopaminergic neurons to ATP10B loss-of-function was also observed in midbrain neuronal cultures derived from ATP10B knock-out human induced pluripotent stem cells clones, which showed a significant reduction in TH-positive neurons.

**Conclusion:** Taken together, our findings demonstrate that ATP10B depletion detrimentally impacts the viability of dopaminergic neurons both *in vivo* and *in vitro*. Moreover, a broader impact on the functionality of the nigrostriatal pathway was evidenced as rats with *ATP10B* knockdown exhibited motor impairments similar to those observed in PD patients.

## Introduction

Parkinson’s disease (PD) stands as the second most prevalent neurodegenerative disorder, with a notable increase in both incidence and prevalence observed over the past two decades. The clinical spectrum of PD is characterized by motor symptoms such as tremor, bradykinesia, rigidity and postural instability, although non-motor aspects also contribute to this disease, such as constipation, cognitive decline, depression, and pain, among others (1). The primary factor behind the motor impairment observed in patients with PD is the gradual neurodegeneration of dopaminergic neurons in the *substantia nigra pars compacta* (*SNpc*) (2). These neurons extend axons to the *dorsal striatum* (*dSTR*) releasing dopamine and establishing the nigrostriatal dopaminergic pathway. This pathway contributes to the basal ganglia circuits involved in action selection, modulation and learning (3,4). Previous research has consistently shown impaired dopamine release from nigrostriatal neurons in numerous models of PD, often preceding or occurring in the absence of neurodegeneration (5). Pathologically, PD is further characterized by the accumulation of aggregated α-synuclein in Lewy bodies and Lewy neurites. These structures, found in the affected brain regions, present a distinctive crowded environment comprising various membrane components, including vesicular structures, dysmorphic organelles like mitochondria, and high lipid content (6). Therefore, the pathophysiology of PD presents a complex interplay between α-synuclein aggregation, dysfunctional mitochondria and lysosomes, disturbed lipid homeostasis and impaired vesicle transport (7,8).

PD presents a multifactorial etiology-involving genetics, environmental factors, and the aging process. In recent years, the PD research community has dedicated substantial efforts to identify novel genetic factors associated with the pathogenesis of this complex disease. Notably, a significant proportion of these newly discovered genes have emerged as key players in the endolysosomal pathway, shedding light on the central role of endolysosomes in the development and progression of PD (7). In a cohort of Belgian patients with PD and dementia with Lewy bodies (DLB) compound heterozygous missense variants were identified in a lysosomal gene from the P4-ATPase family known as *ATP10B*, which was proposed as a novel candidate genetic risk factor in PD. Identified pathogenic variants cause loss of the ATPase activity and are situated in conserved and functionally important domains (9). Although mutations in *ATP10B* have been identified in PD patients from other cohorts, these studies did not confirm that ATP10B rare variants are causally linked to PD or increase the susceptibility to present the disease (10–13). On the other hand, the incomplete penetrance of ATP10B mutations, together with their low frequency and the importance of the *cis/trans* phasing of the compound heterozygous variants, underscore the need to study large cohorts and validate the *cis/trans* configuration to obtain significant results (14–16).

ATP10B is a transmembrane lipid flippase located in late endosomes and lysosomes, coupling lipid export of glucosylceramide (GluCer) and phosphatidylcholine (PC) to ATP hydrolysis (9,17). GluCer is also the main substrate of the lysosomal enzyme glucocerebrosidase (Gcase) encoded by *GBA*. Interestingly, mutations in GBA are one of the main genetic risk factors in PD (18). Lipids are major structural and functional components of cellular membranes. The lipid composition varies not only between membranes of different organelles but also between the cytosolic and exoplasmic leaflets of a single membrane, creating an asymmetric distribution that is a vital characteristic. ATP-dependent flippases play a pivotal role in maintaining membrane asymmetry by facilitating the transbilayer movement of lipids, a process that is energetically unfavorable (19,20). Pathogenic ATP10B variants exhibit a strongly impaired ability to translocate PC and GluCer (9,17). In addition, ATP10B regulates the uptake of PC and affects hexosylceramide levels in human cancer cell models (17). Lysosomal functionality is compromised after *ATP10B* knockdown (KD) in WM-115 melanoma cells, as well as in primary cortical neurons with *ATP10B* KD, where the loss of *ATP10B* also increases susceptibility to cell death. Interestingly, these phenotypes in human cell lines and mouse cortical neurons are exacerbated by exposure to environmental PD risk factors such as rotenone and MnCl_2_ (9).

While *in vitro* models have offered insights into the cellular role of ATP10B, the consequences of ATP10B loss of function on PD neuropathology, particularly related to the function of dopaminergic neurons *in vivo*, remain unclear. To address these questions, we have developed a rat model with targeted *ATP10B* KD in the *SNpc* neurons and investigated its impact on the nigrostriatal dopaminergic pathway.

## Methods

### Study design

Two miRNA-based short-hairpins (miR5 and miR7) targeting distinct regions of rat ATP10B were packaged in an adeno-associated vector (AAV2/7) under a neuronal promoter (CMVenhanced-Synapsin) and unilaterally injected in the right SNpc of adult female Wistar rats. We have previously reported that these two miRNA-based short-hairpins efficiently KD ATP10B in mouse cortical neurons cultures (9). A vector with a scrambled sequence (SCR) was used as control. The AAV2/7 viral vectors were produced by the Leuven Viral Vector Core using the following transfer plasmids for viral vector production: SCR (Addgene plasmid RRID: Addgene_216393), miR5 (Addgene plasmid RRID: Addgene _216391), miR7 (Addgene plasmid RRID: Addgene_216392) (21). The long-term study consisted of 15 to 16 rats per treatment group. Following stereotaxic surgery, behavioral evaluation was performed using tests for spontaneous locomotion (open-field), motor coordination and balance (rotarod), catalepsy (bar test), and motor asymmetry (cylinder, elevated body swing). In addition, in a subset of rats randomly allocated from the initial group (miR5 n=5, miR7 n=5, SCR n=3), *in vivo* striatal dopamine transporter (DAT) binding was performed using ^18^F-FE-PE2I and positron emission tomography imaging (PET). Rats were euthanized 1 year post-injection to investigate neuropathological changes in the brain (miR5 n=8, miR7 n=8, SCR n=8) or changes in protein expression (miR5 n=7, miR7 n=8, SCR n=5). A separate cohort of rats received injections with the same vectors (miR5 n=6, miR7 n=6, SCR n=6) and were sacrificed after 1 month, without behavioral evaluation, to examine possible short-term alterations by immunohistochemistry. An overview of the study design is shown in Fig. 1. Additionally, a smaller group of rats injected with the same vectors (miR5 n=4, miR7 n=4, SCR n=2) and euthanized after 2 weeks was utilized to assess the efficiency of vector transduction in dopaminergic neurons.

**Figure 1.**
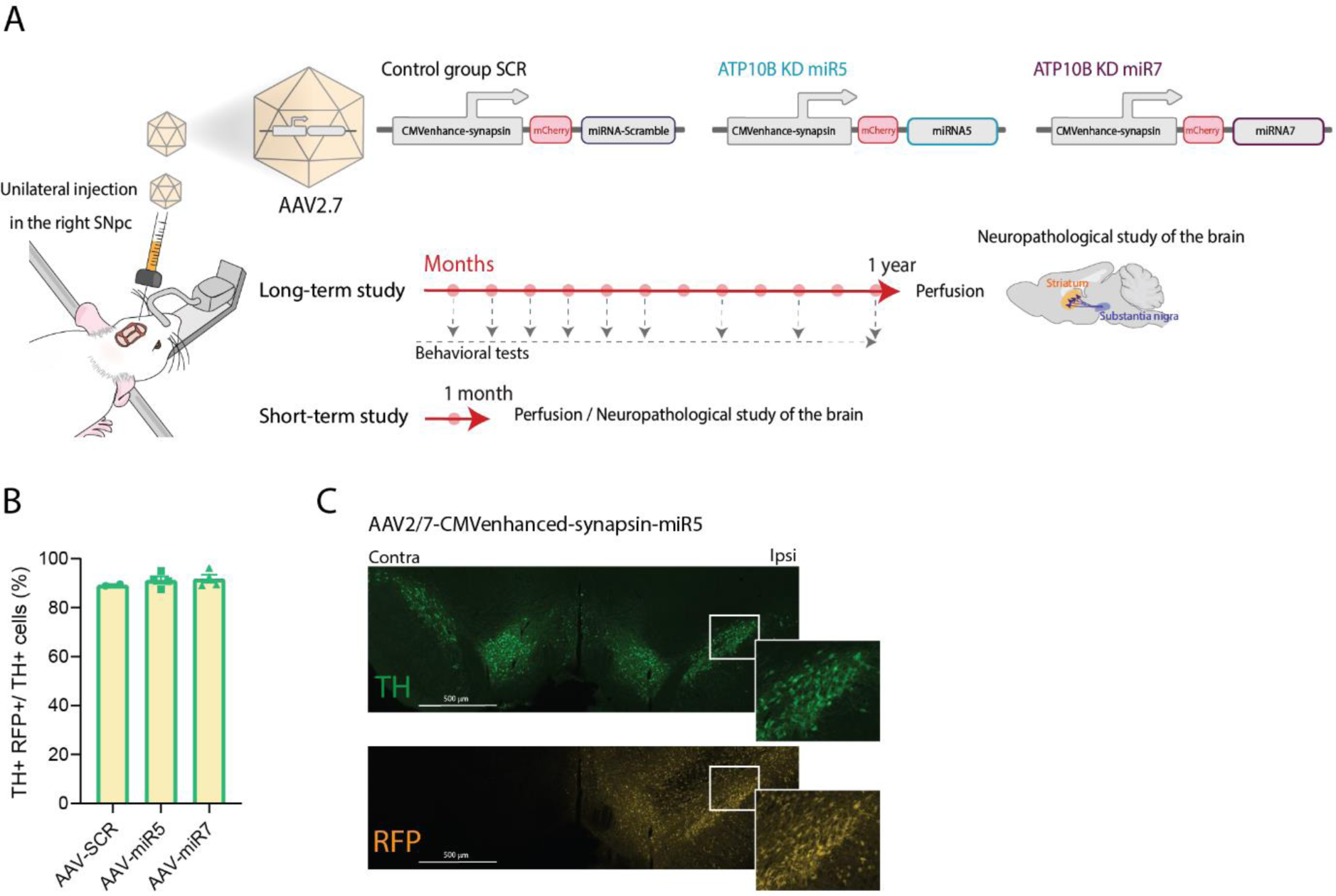
Targeted knockdown of ATP10B in adult rat substantia nigra using AAV vectors. MiRNA-based short-hairpins (miR5 and miR7) targeting distinct regions of ATP10B mRNA, and a scrambled non-targeting sequence (SCR) packaged in AAV2/7 vector under a neuronal promoter (CMVenhanced-synapsin) were stereotactically injected in the right SNpc of rats. (A) Overview of the protocol followed in this study and vector constructs. (B, C) The percentage of double TH+ RFP+ cells within the total TH+ population in the SNpc was determined for the three vectors 2 weeks post-injection. (B) Each data point represents an individual animal, and the number of TH+ and TH+ RFP+ cells from 8 sections was quantified using stereological methods. (C) Representative image of the SNpc of a rat injected with the miR5 vector 2 weeks post-injection, fluorescently stained with TH and RFP antibodies, shows efficient transduction of dopaminergic neurons.

### Animals

Adult (8-9 weeks old) female Wistar rats weighing 200-250 g (Janvier, France) were used for this study, and housed (2-3 animals per cage) in individually ventilated cages with free access to food and water, under a normal 12h-light/12h-dark cycle. Animal experiments were carried out in accordance with the European Communities Council Directive of November 24, 1986 (86/609/EEC) and approved by the Bioethical Committee of the KU Leuven (Belgium) (ECD project 161/2022).

### Stereotaxic surgery

All surgical procedures were performed using aseptic techniques. Rats were anaesthetized with ketamine (60 mg/kg, intraperitoneal (i.p.), Nimatek, Dechra, Belgium) and medetomidine (0.4 mg/kg, i.p., Domitor, Orion Pharma, Finland), and placed in a stereotactic frame (Stoelting, Wood Dale, IL). Rats were injected unilaterally with 3 µL of AAV2/7-CMVenhsynapsin-mCherry-miRSCR, AAV2/7-CMVenhsynapsin-mCherry-miR5, or AAV2/7-CMVenhsynapsin-mCherry-miR7 in the right SNpc (AP: −5.3 mm; L: −2.0 mm; DV: −7.2 mm calculated from dura, using bregma as reference), using a 30-gauge needle and a 10 μl Hamilton syringe (Hamilton, Bonaduz, GR, Switzerland). Each vector was injected at a normalized titer of 7.5 x 10^11^ GC/mL (2.25 x 10^9^ GCs injected per animal), at a flow rate of 0.25 μL/min. Following injection, the syringe was left in place for an additional 5 min, and then slowly retracted. Reversal of anesthesia was performed using atipamezole (0.5 mg/kg, i.p., Antisedan, Orion Pharma) and buprenorphine (0.03 mg/kg, subcutaneous, Vetergesic, Ceva Santé Animale, France) was administered as post-operative analgesia. A detailed description of the protocol can be accessed at protocols.io (dx.doi.org/10.17504/protocols.io.bp2l6b1qzgqe/v1).

### Behavioral tests

Different behavioral tests were performed to assess motor asymmetry and motor function. For each test, rats were acclimatized to the testing room at least 1 h prior to assessment.

#### Rotarod test

Motor coordination and balance was assessed using an accelerated rotarod system (IITC Life Science Rotarod Model I-755, Campden Instruments). Before surgery, rats were trained for 5 min at a constant speed of 5 rpm. During this initial training phase, rats were placed back on the rod after falling. In the second phase of training, rats underwent 3 trials of 1 min at a fixed speed of 5 rpm, 10 rpm, and 15 rpm, with 5 min of rest in-between trials. For testing rotarod performance at baseline, and post-surgery, rats were positioned on rotating rods with a progressively increasing rotation speed, ranging from 4 to 40 rpm over a 5 min period. Three trials were conducted at every timepoint with 5 min resting interval in between. The rotarod protocol was performed on three consecutive days, with the average of the three days taken for statistical analyses. A detailed description of the protocol can be accessed at protocols.io (dx.doi.org/10.17504/protocols.io.5qpvo3zo9v4o/v1).

#### Open field test

The open field set-up consisted of a square box (1 m x 1 m) surrounded by opaque walls that prevent observation of visual cues outside the arena. Rats were placed in the center and spontaneous behavior was recorded using an overhead camera for 5 min. Total distance traveled, velocity, and % immobility time were obtained using AnimalTracker (22), and ipsi-/contralateral 180° turning and rearing frequency were calculated manually in a blinded manner on the acquired videos. An 180° turn was counted if the animal axis performed a spontaneous 180° turn of the body axis while in place. A detailed description of the protocol can be accessed at protocols.io (dx.doi.org/10.17504/protocols.io.kxygx3dozg8j/v1).

#### Cylinder test

The cylinder test was employed to quantify asymmetry in forelimb use. Contacts made by each forepaw with the wall of a 20-cm-wide clear glass cylinder were scored from the videotapes by an observer blinded to the animal’s identity. A minimum of 20 contacts were quantified for each animal. The number of contralateral forelimb contacts was expressed as a percentage of total forelimb contacts. Rats not performing enough contacts were excluded from the analysis. A detailed description of the protocol can be accessed at protocols.io (dx.doi.org/10.17504/protocols.io.kxygx36zdg8j/v1).

#### Elevated body swing test

In the elevated body swing test (EBST), the rat tail was taken approximately 3 cm from its base, and the animal was elevated to about 5 cm above the cage floor and held along the vertical axis for 5 sec. Ipsilateral or contralateral swing were counted manually in a blinded manner, whenever the animal moved its head out of the vertical axis to either side by at least 30° within the 5 s observation interval. The EBST comprised five trials of maximum 5 s, each followed by a brief rest. The mean for all five trials was used for statistical analysis. A detailed description of the protocol can be accessed at protocols.io (dx.doi.org/10.17504/protocols.io.bp2l6x58zlqe/v1).

#### Catalepsy test

The catalepsy bar test is used to measure the failure to correct an imposed posture resulting from muscular rigidity. For this test, the forepaws of the rats were placed on an elevated bar with the hind paws remaining on the floor. The time for the rat to correct this posture was recorded and used an index of the intensity of catalepsy. Three trials were performed, with a brief rest in between. The average of all three trials was used for statistical analysis. A detailed description of the protocol can be accessed at protocols.io (dx.doi.org/10.17504/protocols.io.q26g7p24kgwz/v1).

### Small animal DAT microPET imaging

For longitudinal *in vivo* measurement of DAT integrity, we used ^18^F-FE-PE2I, a highly selective DAT PET radioligand (23), ^18^F-FE-PE2I produced under GMP was obtained from the hospital radiopharmacy at UZ Leuven.

Rats were anesthetized (5% isoflurane in O_2_ for induction and 2% thereafter at 1 L/min flow rate), placed on a heated mat and cannulated with a 23G catheter. They were then placed on the imaging bed (Molecubes, Ghent, Belgium) with integral temperature and respiration monitoring before being transferred to a β-cube microPET scanner (Molecubes/Bruker, Ghent, Belgium). Dynamic PET images were acquired for 90 min starting from intravenous injection with ^18^F-FE-PE2I (8.9±2.7 MBq)/rat. Specific activity was 286±119 GBq/micromol with mass doses injected between 0.1─0.4 nmol. Rats were kept under anesthesia during the entire procedure (2.5% isoflurane in O_2_ at 1 L/min flow rate), with temperature and respiration monitored throughout. After PET scanning, a computed tomography (CT) image was acquired for anatomic coregistration with an X-cube CT scanner (Molecubes, Ghent, Belgium), using the following parameters: 50kVp, 480 exposures, 85 ms/projection, 100 μA tube current, rotation time 60 s. After scanning, rats were recovered, with on average two weeks between scanning sessions.

### Image processing and analysis

PET list mode data were binned into 16 frames (4 × 15 seconds, 4 × 1 min; /1 x 5 min /5 x 10 min and 2 x 15 min) and reconstructed into a 192 × 192 image matrix with 0.4 mm voxel size, using 30 iterations using the native Maximum-Likelihood Expectation-Maximization (MLEM) algorithm. Correction was done for randoms, scatter, attenuation based on CT, and data were scaled to kBq/cc after calibration to a standard ^18^F-phantom. Images were decay-corrected to the start of the scan. CT data were reconstructed using a regularized iterative algorithm (24) with a voxel size of 200 µm (isotropic) and were scaled to Hounsfield Units (HUs) after calibration against a standard air/water phantom. PET and CT data were then cropped to the skull and CT scans were co-registered to an in-house CT skull template using an affine transformation. The first 9 frames of PET data were summed for rigid co-registration with the CT scan. Dynamic PET data were then transformed to template space using the CT-based transformation matrices. Striatal and cerebellar regions of interest in template space (25) and time-activity curves were created (operations carried out using PFUS). Time activity curves were imported in PKIN v4.0 (PMOD Technologies GmbH, Zürich, Switzerland), where a standard reference tissue model (SRTM) was applied to generate non-displaceable binding potential (BPnd) values as measure for absolute tracer binding, using the cerebellum as the reference tissue (26). Parametric BPnd images were then generated using the voxel SRTM method in PXMOD (PMOD Technologies GmbH, Zürich, Switzerland) with the same reference region.

### Immunohistochemical staining

For the staining of the brain, rats were sacrificed with an overdose of sodium pentobarbital (200 mg/kg, i.p., Dolethal, Vetoquinol, Lure, France), and transcardially perfused with saline and 4% paraformaldehyde (PFA) in PBS. After post-fixation overnight in 4% PFA, 20 µm thick coronal brain sections were made with a vibrating microtome (HM 650 V, Microm).

Immunohistochemistry of the STR was performed on free-floating sections. Sections were washed with PBS, and incubated in an antigen retrieval solution (0.1 M citrate buffer pH 6.0) for 30 min at 80°C. After 20 min on ice, the sections were washed in PBS, and incubated in 3% H_2_O_2_ and 10% methanol in PBS for 10 min at room temperature. The sections were washed 2 x 5 min in PBS with 0.1% Tergitol (PBS-T), and incubated overnight in primary antibody against tyrosine hydroxylase (TH) 1:10.000 (rabbit anti-TH, Millipore Cat# AB152, RRID:AB_390204), diluted in PBS-T with 10% goat serum at room temperature. The next day, the sections were washed 2 x 5 min in PBS-T, followed by secondary antibody incubation with biotinylated anti-rabbit IgG (DakoCytomation). Then, the sections were washed 2 x 5 min in PBS-T and incubated with streptavidin–horseradish peroxidase complex (DakoCytomation). Following 2 x 5 min washes in PBS-T, TH immunoreactivity was visualized using DAB 3,3’diaminobenzidine tetrahydrodhloride (D5905, Sigma). A description of the protocol can be accessed at protocols.io (dx.doi.org/10.17504/protocols.io.eq2lypnkqlx9/v1).

Images of dSTR sections were captured with the Aperio CS2 slide scanner. The QuPath software was employed to measure the percentage TH positive area in seven sections per animal across the dSTR. Every twelve sections throughout the entire dSTR was analyzed, with a total of 8 sections for each animal.

TH Immunofluorescence of SNpc was performed on sections mounted on Superfrost plus glass slides, dried overnight and following an antibody signal enhancement (ASE) protocol (27). Sections were pretreated with antigen retrieval solution (Tris-HCl-EDTA buffer pH 9.0 + 0.05% SDS) in the steamer for 30 min. After 20 min on ice, sections were washed with ASE wash buffer (PBS + 0.5% Tween-20) and blocked for 30 min with ASE blocking solution (PBS + 2% donkey serum, 50 mM glycine, 0.05% Tween-20, 0.1% Tergitol, 0.1% BSA). Sections were incubated overnight at 4°C with primary antibody TH 1:1000 (TH, Aves Labs Cat# TYH, RRID:AB_10013440) in ASE primary antibody buffer (PBS + 10 mM glycine, 0.05% Tween-20, 0.1% Tergitol, 0.1% H_2_O_2_). Next day, after one rinse and 2 x 3 min washes with ASE wash buffer, sections were incubated for 2 h at room temperature with secondary antibody donkey anti-chicken Alexa 647 1:500 diluted in ASE secondary buffer (PBS + 0.1% Tween-20). After being rinsed in PBS and allowed to dry, the sections were covered with Mowiol (Calbiochem, San Diego, CA). A description of the protocol can be accessed at protocols.io (dx.doi.org/10.17504/protocols.io.n2bvj37zplk5/v1).

For evaluating transduction efficiency and performing lysosomal staining of nigral dopaminergic neurons, sections of the SNpc were pretreated with antigen retrieval solution (0.1 M citrate buffer pH 6.0) for 30 min at 80°C. After 20 min on ice, sections were washed with PBS and blocked for 1 h with blocking solution (PBS-T + 10% donkey serum). Sections were incubated overnight at room temperature with primary antibodies against TH 1:1000 (chicken anti-TH, Aves Labs Cat# TYH, RRID:AB_10013440), RFP 1:1000 (rabbit anti-RFP, Rockland Cat# 600-401-379S, RRID:AB_11182807), LAMP1 1:500 (rat anti-LAMP1 clone 1D4B, Novus Cat# NBP1-49151, RRID:AB_10011343), LAMP2a 1:500 (rat anti-LAMP2a, Abcam Cat# ab13524, RRID:AB_2134736), GCase 1:500 (rabbit anti-GBA C-terminal, Sigma-Aldrich Cat# G4171, RRID:AB_1078958) and Cathepsin B 1:500 (goat anti-Cathepsin B, R&D Systems Cat# AF965, RRID:AB_2086949) in primary antibody buffer (PBS-T + 10% donkey serum). Next day, after one rinse and 2 x 5 min washes with PBS-T, sections were incubated for 2 h at room temperature with secondary antibody donkey anti-chicken Alexa 488 or 555 1:500 (for TH), donkey anti-rabbit Alexa 555 1:500 (for RFP), donkey anti-rabbit Alexa 647 1:500 (for GCase), donkey anti-goat Alexa 647 1:500 (for Cathepsin B) and donkey anti-rat Alexa 488 1:500 (for LAMP1 and LAMP2a), diluted in PBS-T. After being rinsed in PBS and allowed to dry, the sections were coverslipped with Mowiol (Calbiochem, San Diego, CA). A description of the protocol can be accessed at protocols.io (dx.doi.org/10.17504/protocols.io.3byl4qoxrvo5/v1).

The number of TH+ cells in the SNpc (1 month and 1 year time points) or the number of TH+/RFP+ cells (2 week time point) was determined by stereological measurements using the optical fractionator method in a computerized system as described before, in a blinded manner (Stereo Investigator; MicroBrightField (MBF), Delft, The Netherlands). Every ten sections throughout the entire SNpc was analyzed, with a total of 7 sections for each animal.

The number and volume of the late endosomes/lysosomes was determined in TH+ dopaminergic neurons in three sections per animal spanning the SNpc. Images were captured using a Nikon AX confocal microscope at 60x magnification. The captured Z-stacks were 3D reconstructed using Imaris version 10.0.1 (Oxford Instruments, Abingdon, United Kingdom), and labeled (LAMP1+, LAMP2a+, Cathepsin B+, and GCase+) organelles were quantified using Imaris Spot detection within delineated TH+ cell surfaces within the field of view.

### Protein extraction and Western blot

For protein expression analysis, rats were sacrificed with an overdose of sodium pentobarbital (200 mg/kg, i.p., Dolethal, Vetoquinol, Lure, France), and transcardially perfused with saline. SN tissue was freshly isolated, snap-frozen and stored at -80°C until analysis. Samples were weighed and homogenized in 10 volumes of RIPA buffer (50 mM Tris–HCl, 150 mM NaCl, 0.1% (w/v) SDS, 1% (v/v) Triton-X100, 0.5% (w/v) Sodium Deoxycholate, 1.0 mM EDTA pH 7.4) containing a protease inhibitor cocktail (Roche cOmplete) and phospho-STOP EASYPACK (Roche) using a tissue homogenizer (TH, Omni Tissue Homogenizer). After homogenization, samples were sonicated 3 times during 15 sec, and centrifuged at 6000 g for 10 min at 4 °C. Protein sample concentration was determined by BCA protein assay (Thermo Scientific, MA, USA) according to the manufacturer’s directions.

To assess LAMP1, GCase, Cathepsin B, p62 and α-synuclein protein levels, Western blots were performed using 15 μg of SN extracts prepared in 4x Leammli buffer (0.24M Tris pH 6.8, 7.27% SDS, 40% Glycerol, 10% β-Mercaptoetanol, 0.01% Bromophenol blue) and boiled for 10 min at 98°C. Protein samples were loaded on 4–15% Criterion™ Tris-HCl Protein Gel and transferred to a polyvinylidene fluoride (PVDF) membrane (Bio-Rad). Nonspecific binding sites were blocked for 1 h at room temperature in 5% nonfat milk in PBS-T. After overnight incubation at 4°C with primary antibodies (LAMP1, Abcam Cat# ab25630, RRID:AB_470708 1:1000, GCase Sigma-Aldrich Cat# G4171, RRID:AB_1078958 1:1000, Cathepsin B R&D Systems Cat# AF965, RRID:AB_2086949 1:1000, P62/SQTM1 Proteintech Cat# 55274-1-AP, RRID:AB_11182278 1:1000, α-synuclein BD Biosciences Cat# 610786, RRID:AB_398107 1:1000, βactin Sigma-Aldrich Cat# A5441, RRID:AB_476744 Sigma 1:5000) blots were washed 3 times during 10 min with PBS-T and incubated with horseradish peroxidase-conjugated secondary antibody (Dako) for 1 h and washed again 3 times. Bands were visualised using Clarity Western ECL (Bio-Rad) and developed with a GE ImageQuant 800 (GE Healthcare). Densitometric analysis was performed using ImageQuant software (GE Healthcare).

### Generation of isogenic knockout lines

The human induced pluripotent stem cells (iPSCs) BJ-SiPS line was cultured using StemflexTM medium (ThermoFisher). Before nucleofection, the cells were pre-treated with a 10 μM Rhok inhibitor for an hour. The cells were then dissociated using Accutase, after which they were pelleted and resuspended in 800 μl PBS containing 5 μg of px330 CRISPR DNA each. Finally, they were transferred into nucleofector cuvettes. Nucleofection was performed using the P4 Nucleofector kit from Amaxa and the standard and program hiPSC CA-137. Two CRISPR sgRNA targeting exon1 were used to produce the knockout (KO) (CR1: CACCGCTACAACTTGACACAGCAG, AAACCTGCTGTGTCAAGTTGTAGC CR2: CACCGAATTGCTCAAAGAGATTCCG, AAACCGGAATCTCTTTGAGCAATTC), genotype primers used were F: TGGCAGTGGAGAGTCAGAGA, R: CCTGGGGAACAGAATGAGAC. Clones with homozygous or compound heterozygous deletions, leading to truncations and frameshift mutations, were identified using genotyping PCR. Clones for all lines containing deletions were identified by Sanger sequencing. A description of the protocol can be accessed at protocols.io (dx.doi.org/10.17504/protocols.io.bu7znzp6)

### Midbrain differentiation

Human induced pluripotent stem cells (hiPSCs) were cultured in StemflexTM medium (ThermoFisher) at 37°C, with 5% CO_2_ in a humidified incubator, as previously described. Differentiation was done by dual-SMAD inhibition with SB431542 (R&D Systems, 10 μM), LDN193189 (Stemgent, 100 nM), B27 minus Vit A and N2 in DMEM-F12. Midbrain-specific patterning for midbrain NPCs was made with the addition of CHIR99021 (Stemgent, 3 μM), Purmorphamine (STEMCELL, 2 μM), and SAG (Abcam, 1 μM). Post patterning Neural maturation medium was DMEM F12 medium containing N2, B27-VitA, 20 ng/mL GDNF (R&D Systems), 20 ng/mL BDNF (R&D Systems), 0.2 mM ascorbic acid (Sigma), 0.1 mM dibutyryl cAMP (Biolong), 10 μM DAPT (Cayman Chemical). The medium for long-term culture was DMEM F12 medium containing N2, B27-VitA, 10 ng/ML GDNF (R&D Systems), 10 ng/mL BDNF (R&D Systems), 0.2 mM ascorbic acid (Sigma). A description of the protocol can be accessed at protocols.io (dx.doi.org/10.17504/protocols.io.bp2l62dj5gqe/v1)

### Staining and quantification

Cells were fixed with 4% PFA for 20 min, blocked in 0.1% Triton X-100 in 5% horse serum/PBS, and then incubated in primary antibodies overnight at 4°C (TH: Millipore Cat#MAB318; RRID: AB_2201528, MAP2: RRID: AB_2564858) The following day, cells were washed and incubated in secondary antibodies (Donkey anti-Mouse IgG, Alexa Fluor 555 Invitrogen Cat#A-31570, Goat anti-Chicken IgY (H+L) Alexa Fluor 488 RRID: AB_2534096) and DAPI nuclear stain according to protocol. Imaging was performed using the High-content imager CX7 with phenotyping and quantifications using CellProfiler. The average of at least three fields of a well was counted as the N value for statistical analysis. A detailed description of the protocol can be accessed at protocols.io. (dx.doi.org/10.17504/protocols.io.j8nlk8wowl5r/v1)

### Statistical analysis

Data are presented as mean ± standard error of the mean (s.e.m.), and individual values. Statistical analyses were performed using GraphPad Prism 10.2.0. For analyzing paired observations within one group of animals, we used a paired t-test. For all the other analyses, we employed a one-way ANOVA, followed by post-hoc comparisons. Parametric or non-parametric statistical tests were performed based on the normality of residuals and equal variance across groups, tested using the D’Agostino-Pearson omnibus and Brown-Forsythe tests respectively. The α-value was set at 0.05.

## Results

### *In vivo* KD of ATP10B in adult rat SN by stereotactic injection of AAV vectors

*ATP10B* has recently emerged as a candidate genetic risk factor in the context of PD. Studies have identified loss of function mutations and reduced ATP10B mRNA expression in PD patients (9). To gain a deeper understanding of ATP10B’s role in PD pathogenesis, and specifically in the nigrostriatal dopaminergic system, we conducted a study employing a rat model with ATP10B KD targeted to the SNpc neurons (Fig. 1A). To knock down *ATP10B*, we used two different miRNA-based shRNA sequences (miR5 and miR7) that we previously showed to induce loss of ATP10B expression and sensitize primary mouse cortical neuron cultures of PD-related stressors (9). A scrambled sequence (SCR) was used as control. Two weeks post-injection, we assessed the transduction efficiency and construct expression of the AAV2/7 miR5, miR7, and SCR vectors in SNpc dopaminergic neurons based on mCherry expression. Our findings revealed that all three vectors performed similarly, demonstrating high transduction rates of TH+ cells (∼90%) (Fig. 1B, C).

### ATP10B KD in the SNpc neurons leads to decreased motor function and induces motor asymmetry at 12 months post injection

To evaluate the motor performance of the ATP10B KD rats, we conducted a series of behavioral tests at different times post injection. We will focus on the behavioral test results at 1 year, as this time point was chosen for the neuropathological examination of the brain, although motor impairments could be observed from earlier time points post injection (Fig. S1). In the rotarod test, no significant differences could be observed between the groups at baseline, indicating equal acquisition of rotarod motor skill (data not shown). At 12 months post-injection, we noted an overall compromised motor coordination and balance, as evidenced by the time spent on top of the rod across the three groups (treatment factor: p = 0.0083). Post hoc analysis revealed that a decreased time on top of the rod was observed in the miR5 group compared to the SCR (miR5 vs. SCR: p = 0.0043) (Fig. 2A). An overall treatment effect was also evidenced in the cylinder test, where the percentage of left (contralateral) forepaw usage was quantified (treatment factor: p = 0.0110). Post hoc analysis revealed that miR5 group had a significant decrease in left forepaw usage when compared to SCR (miR5 vs. SCR: p = 0.0068) (Fig. 2B). This motor asymmetry observed in the cylinder test was consistent with the findings in the EBST (Fig. 2C) and the turning behavior in the open field (Fig. 2H). Indeed, both the miR5 and miR7 groups exhibited a strong preference for performing ipsilateral body swings in the EBST (treatment factor: p = 0.0016; miR5 vs. SCR: p = 0.0321, miR7 vs. SCR: p = 0.0009) (Fig. 2C) and spontaneous ipsilateral rotations in the open field (treatment factor: p = 0.0123; miR5 vs. SCR: p = 0.0388, miR7 vs. SCR: p = 0.0112) (Fig. 2H) compared to the SCR group. In addition, overall impaired locomotor activity and exploration behavior was also observed in the open field, indicated by parameters such as distance traveled (treatment factor: p = 0.0433), velocity (treatment factor: p = 0.0487) and frequency of rearing (treatment factor: p = 0.0064) (Fig. 2E-G). While post hoc analysis revealed that the miR7 group exhibited a more pronounced impairment in distance traveled (miR7 vs. SCR: p = 0.0312) and velocity (miR7 vs. SCR: p = 0.0212), both miR5 and miR7 groups decreased their rearing behavior in the open field (miR5 vs. SCR: p = 0.083, miR7 vs. SCR: p = 0.0166). Notably, the catalepsy test evidenced an increased muscular rigidity in the miR5 and miR7 groups compared to SCR control group (treatment factor: p = 0.0174; miR5 vs. SCR: p = 0.0181, miR7 vs. SCR: p = 0.0481). This was indicated by a more prolonged duration of holding the elevated bar before correcting their posture to the floor (Fig. 2D). In summary, ATP10B KD in the SNpc neurons using 2 different short hairpin sequences leads to a spectrum of behavioral impairments, encompassing motor asymmetry and dysfunction, similar to those observed in PD patients.

**Figure 2.**
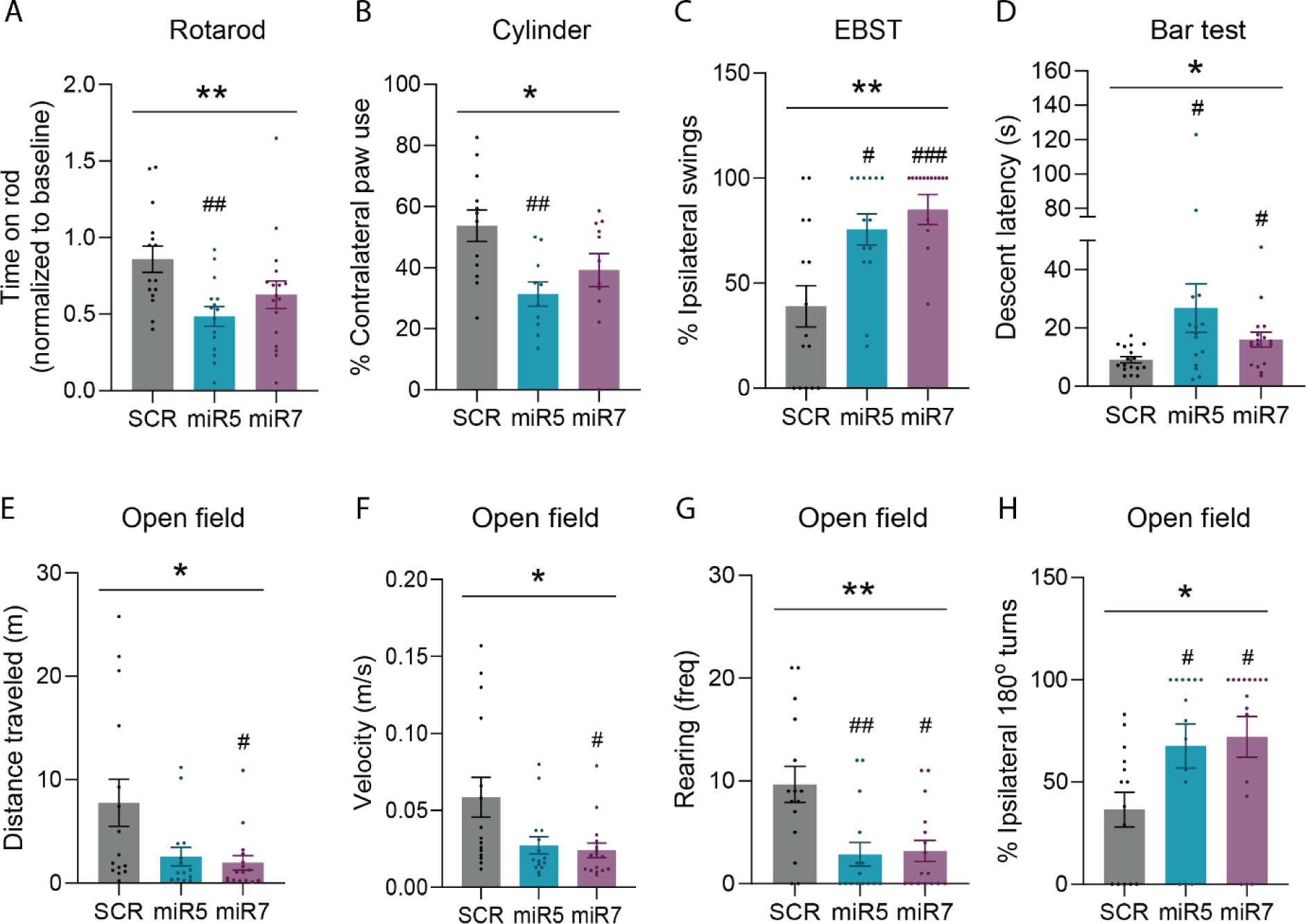
**ATP10B KD in SNpc neurons leads to Parkinsonian motor deficits at 12 months post injection**. (A) Motor coordination and balance was assessed using an accelerated rotarod test (4-40 rpm), with the average performance post-lesion normalized to the individual rat performance prior to surgery (p = 0.0083; SCR vs. miR5 p = 0.0043, SCR vs. miR7 p = 0.1111). (B, C, H) Motor asymmetry was examined by quantifying the preference in the use of the contralateral paw in the cylinder test (p = 0.0110; SCR vs. miR5 p = 0. 00068, SCR vs. miR7 p = 0. 0799), the bias to ipsilateral swing in the EBST (p = 0.0016; SCR vs. miR5 p = 0.0321, SCR vs. miR7 p = 0.0009), and the spontaneous ipsilateral turning behavior in the open field test (p = 0.0123; SCR vs. miR5 p = 0.0388, SCR vs. miR7 p = 0.0112). (E, F, G) Spontaneous motor behavior was recorded during a 5-min open field test and analyzed for total distance traveled (p = 0.0380; SCR vs. miR5 p = 0.1035, SCR vs. miR7 p = 0.0312), velocity (p = 0.0249; SCR vs. miR5 p = 0.0722, SCR vs. miR7 p = 0.0212), and frequency of rearing (p = 0.0064; SCR vs. miR5 p = 0.0083, SCR vs. miR7 p = 0.0166). (D) The catalepsy bar test was performed to assess muscular rigidity. Time spent by the rat grabbing the elevated bar before correcting its posture on the floor was measured (p = 0.0174; SCR vs. miR5 p = 0.0181, SCR vs. miR7 p = 0.0481). In (A, C-H) data are mean ± s.e.m, and analyzed using non-parametric one-way ANOVA (Kruskall Wallis) and Dunn’s post-hoc test versus SCR. SCR (n=15), miR5 (n=15), miR7 (n=17). In (B) data are mean ± s.e.m and analyzed using one-way ANOVA and Dunnet’s post-hoc test versus SCR. SCR (n=12), miR5 (n=10), miR7 (n=11). Each dot represents the value of one animal.

### ATP10B KD in the SNpc neurons results in a progressive loss of striatal DAT in the presynaptic terminals of dopaminergic neurons

Longitudinal imaging with ^18^F-FE-PE2I was done on miR5 and SCR rats at four distinct time points: 2-4 months, 4-6 months, 7-9 months, and 12 months post-injection. The results are expressed as right versus left BPnd of ^18^F-FE-PE2I in the STR of miR5 and SCR rats. BPnd values obtained from both hemispheres for the different groups and the different time points are shown in the Fig. S2. While SCR rats consistently exhibited a stable BPnd on both sides of the STR over time, ATP10B KD miR5 rats displayed a reduction in ^8^F-FE-PE2I BPnd on the ipsilateral side (right) compared to the contralateral side (left) (Fig. 3A, B). Additionally, the right versus left BPnd value in this group decreased progressively over time (2-4 months: ∼13% p = 0.013, 4-6 months: ∼21% p = 0.045, 7-9 months: ∼26% p = 0.0055, 12 months: ∼34% p = 0.0024), indicating a gradual loss of DAT in the presynaptic terminals of dopaminergic neurons. Similar directions of change were observed in a smaller sample of ATP10B KD miR7 rats imaged at 7-9 months (∼66%) and 12 months (∼53%) post injection (Fig. 3A, B).

**Figure 3.**
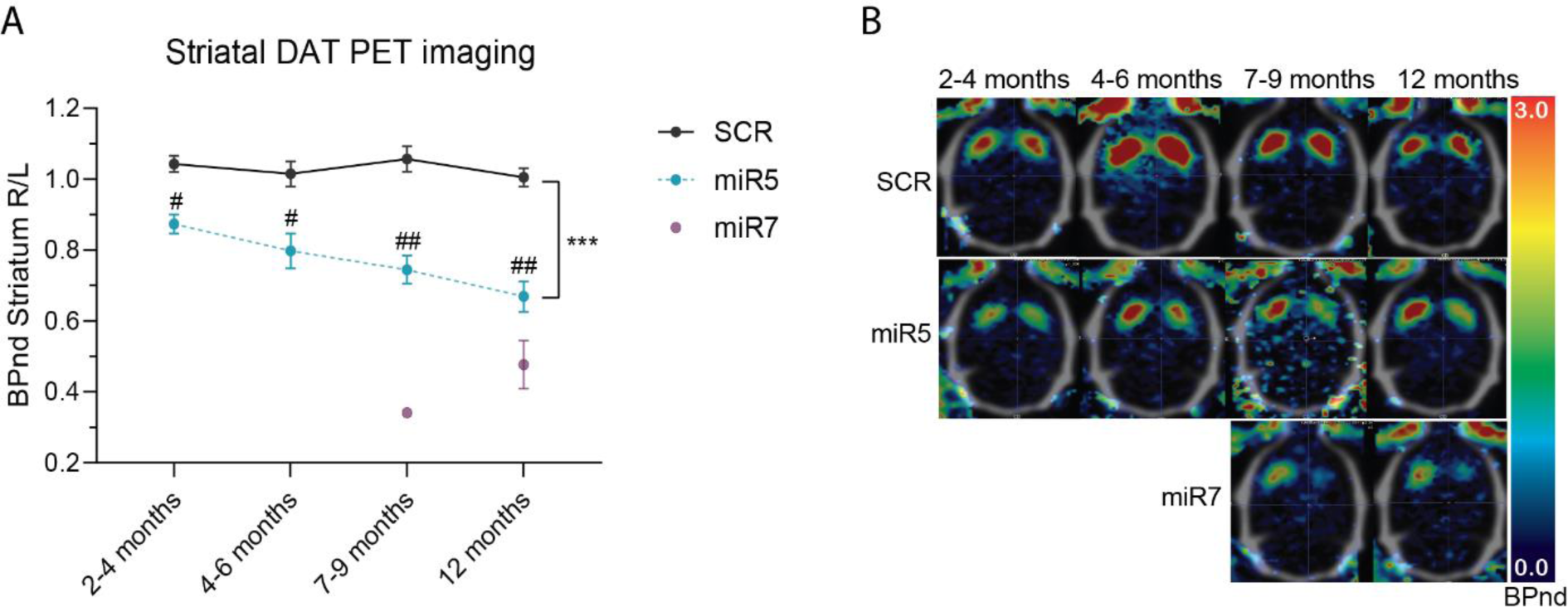
ATP10B KD in SNpc neurons leads to decreased striatal dopamine transporter (DAT) binding. (A) Striatal DAT binding potential (BPnd), measured by ^18^F-FE-PE2I microPET imaging, in the ipsilateral (R) vs. the contralateral (L) striatum in miR5-injected animals (n=5) and SCR-injected animals (n=3) (treatment factor p = 0.0007; SCR vs. miR5 2-4 months p = 0.0134, SCR vs. miR5 4-6 months p = 0.0448, SCR vs. miR5 7-9 months p = 0.0448, SCR vs. miR5 12 months p = 0.0024). Data are mean ± s.e.m. and analyzed using two-way ANOVA and Sidak’s post hoc test miR5 vs. SCR at corresponding time point. The effect could be reproduced in miR7-injected animals at 7-9 months (n=2) and 12 months (n=3). (B) Parametric images of ^18^F-FE-PE2I BPnd in representative SCR (upper) and miR5 (middle) rats over time; lower row represents representative miR7 images at the final two time points. Transverse images are shown in neurological orientation and are overlaid onto a reference CT skull template with the miR5/miR7 lesioned striatum on the right-hand side.

### ATP10B KD in the SNpc results in a progressive decrease of dopaminergic terminals in the striatum

At 1 month post-injection, we observed a significant yet subtle decrease in dopaminergic terminals in the miR5 group, indicated by a reduction in TH+ area in the ipsilateral compared to the contralateral dSTR (miR5 ipsi vs. contra: p = 0.0046) (Fig. 4A, B). At one year post-injection there was a significant decline in dopaminergic terminals when comparing the ipsilateral vs. contralateral dSTR in both miR5 (∼25%) and miR7 (∼42%) rats, which was not present in the SCR group (miR5 ipsi vs. contra: p = 0.0025, miR7 ipsi vs. contra: p = 0.0026) (Fig. 4C, D). An initial pilot study in a separate cohort of rats injected with AAV2/7 SCR and miR5 vectors showed comparable results at 1 year post injection, confirming the impact of ATP10B on dopaminergic terminal loss (Fig. S3). In conclusion, the depletion of ATP10B in the neurons of the SNpc in rats results in a gradual loss of dopaminergic terminals in the dSTR.

**Figure 4.**
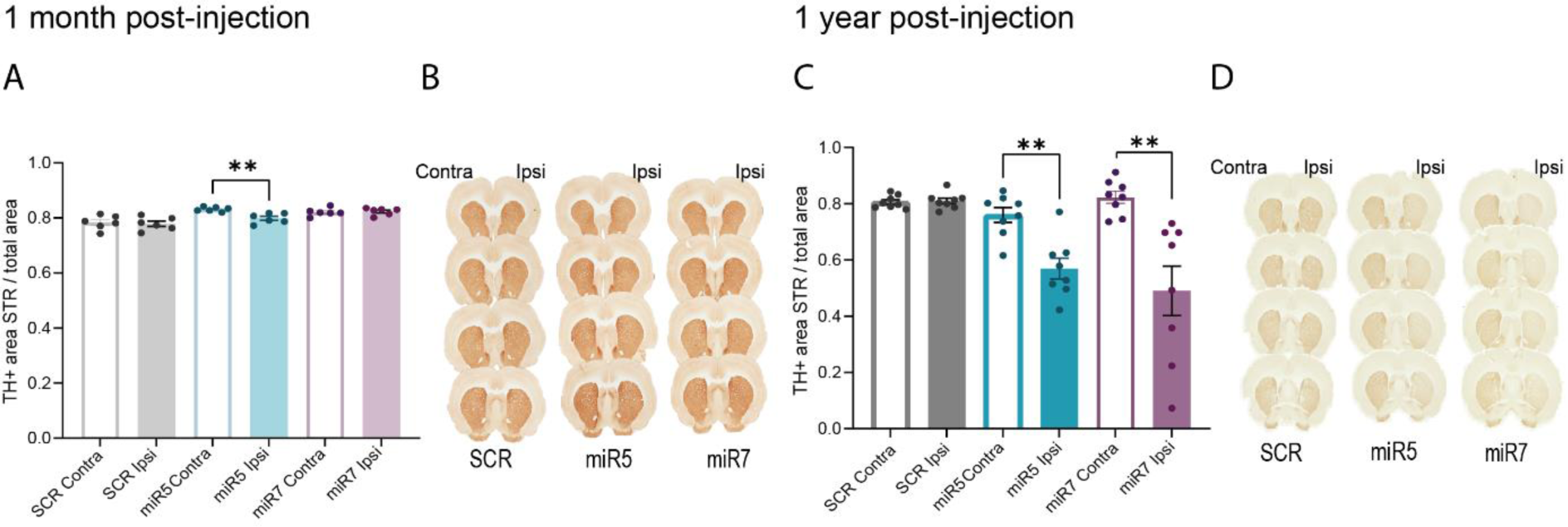
**ATP10B KD in SNpc neurons leads to time-dependent loss of dopaminergic terminals in the dSTR**. TH positive area was quantified using QuPath in 8 different sections that cover the STR. (A) TH positive area in the contralateral (Contra) and ipsilateral (Ipsi) dorsal STR of 1 month post-injection rats (SCR Contra vs. SCR Ipsi p = 0.5704, miR Contra vs. miR5 Ipsi p = 0.0046, miR7 Contra vs. miR7 Ipsi p = 0.6951). SCR (n=6), miR5 (n=6), miR7 (n=6). (B) Representative image of TH immunohistochemical staining on 4 different sections from one SCR, miR5 and miR7 rat 1 month post-injection. (C) TH positive area in the contralateral and ipsilateral dorsal STR of 1 year post-injection rats(SCR Contra vs. SCR Ipsi p = 0.5397, miR Contra vs. miR5 Ipsi p = 0.0025, miR7 Contra vs. miR7 Ipsi p = 0.0026). SCR (n=8), miR5 (n=8), miR7 (n=8). (D) Representative image of TH immunohistochemical staining on 4 different sections from one SCR, miR5 and miR7 rat 1 year post-injection. Data are mean ± s.e.m, and anayzed using a paired t-test. Each dot represents the average of the 8 sections per animal.

### ATP10B KD in the SNpc results in a progressive loss of dopaminergic neurons and altered expression of endolysosomal markers in surviving dopaminergic neurons

At 1 month post injection, a subtle but significant reduction in the number of TH+ dopaminergic neurons was observed in the ipsilateral vs. contralateral SNpc of miR7 rats (miR7 ipsi vs. contra: p = 0.0160) (Fig. 5A, B). At 1 year post injection, both the miR5 and miR7 groups, but not the SCR group, exhibited a decrease in the number of TH+ dopaminergic neurons on the ipsilateral compared to the contralateral SNpc (miR5 ipsi vs. contra: p = 0.0071, miR7 ipsi vs. contra: p < 0.0001) (Fig. 5C, D).

**Figure 5.**
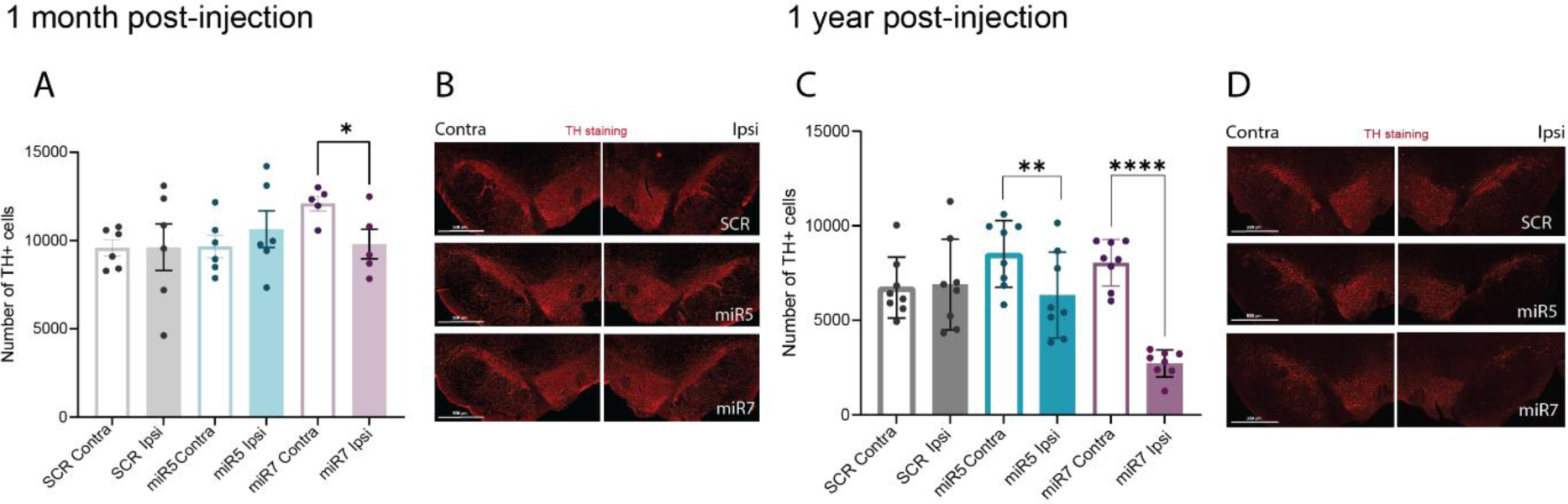
ATP10B KD in SNpc neurons leads to time-dependent loss of dopaminergic neurons in the SNpc. TH positive cells were counted using stereology in 7 different sections that cover the SNpc. (A) Number of TH+ cells in the contralateral (Contra) and ipsilateral (Ipsi) SNpc of 1 month post-injection rats(SCR Contra vs. SCR Ipsi p = 0.98, miR Contra vs. miR5 Ipsi p = 0.2227, miR7 Contra vs. miR7 Ipsi p = 0.0160). SCR (n=6), miR5 (n=6), miR7 (n=5). (B) Representative image of TH immunofluorescence staining on 1 SNpc section from one SCR, miR5 and miR7 rat 1 month post-injection. (C) Number of TH+ cells in the contralateral (Non Inj) and ipsilateral (Inj) SNpc of 1 year post-injection rats(SCR Contra vs. SCR Ipsi p = 0.7125, miR Contra vs. miR5 Ipsi p = 0.0071, miR7 Contra vs. miR7 Ipsi p < 0.0001). SCR (n=8), miR5 (n=8), miR7 (n=8). (D) Representative image of TH immunofluorescence staining on 1 SNpc section from one SCR, miR5 and miR7 rat 1 year post-injection. Data are mean ± s.e.m, and analyzed using a paired t-test. Each dot represents the average of the 7 sections per animal.

Previous *in vitro* studies have demonstrated that loss of ATP10B has an impact on lysosomal activity. To investigate this further in our model, we first performed western blot analysis to assess the total levels of various lysosomal proteins in whole SN extracts obtained from miR5, miR7 and SCR groups at 12 months post-surgery. Specifically, we quantified the expression of LAMP1, GCase and Cathepsin B, as well as the autophagy receptor p62. Given the importance of lysosomes and autophagy in the clearance and degradation of α-synuclein, we also measured total α-synuclein levels. However, no global differences were observed between ATP10B KD and SCR groups for any of these proteins in total SN extracts (Fig. S4).

Next, we wanted to further explore lysosomal changes that may occur specifically in SNpc dopaminergic neurons and might be masked in whole tissue extracts. Consequently, we conducted immunofluorescence staining of different late endosomal/lysosomal markers together with TH in the SNpc of the ATP10B KD rats (Fig. 6A). Interestingly, the miR7 group presented a decrease in the number of LAMP1+ and LAMP2a+ late endosomes/lysosomes in dopaminergic neurons compared to SCR (LAMP1 miR7 vs. SCR: p < 0.0001, LAMP2a miR7 vs. SCR: p < 0.0001). Additionally, the LAMP1+ and LAMP2a+ late endosomes/lysosomes in the miR7 injected animals displayed an enlarged volume (LAMP1 miR7 vs. SCR: p = 0.006, LAMP2a miR7 vs. SCR: p = 0.0009) (Fig. 6B, C). When examining the expression of the lysosomal hydrolase cathepsin B, a reduction was noted in the number of cathepsin B+ organelles in the dopaminergic neurons of both the miR7 and miR5 groups compared to SCR (miR7 vs. SCR: p < 0.0001, miR5 vs. SCR: p = 0.0016). Notably, these lysosomes also presented an enlarged volume, an effect particularly noticeable in miR7 injected rats (miR7 vs. SCR: p < 0.0001) (Fig. 6D). Although no alterations were detected in the number of Gcase+ organelles in any of the KD groups, a slight increase in Gcase+ lysosomal volume was discerned in both the miR5 and miR7 groups compared to SCR (miR5 vs. SCR: p = 0.0012, miR7 vs. SCR: p = 0.0020) (Fig. 6E). It is worth noting that rats injected with the control SCR vector showed an increase in the number of late endosomes/lysosomes in the dopaminergic neurons of the injected side compared to the non-injected side (Fig. 6B-D), suggesting an effect attributed to the injection of a viral vector containing a shRNA sequence and RFP expression.

**Figure 6.**
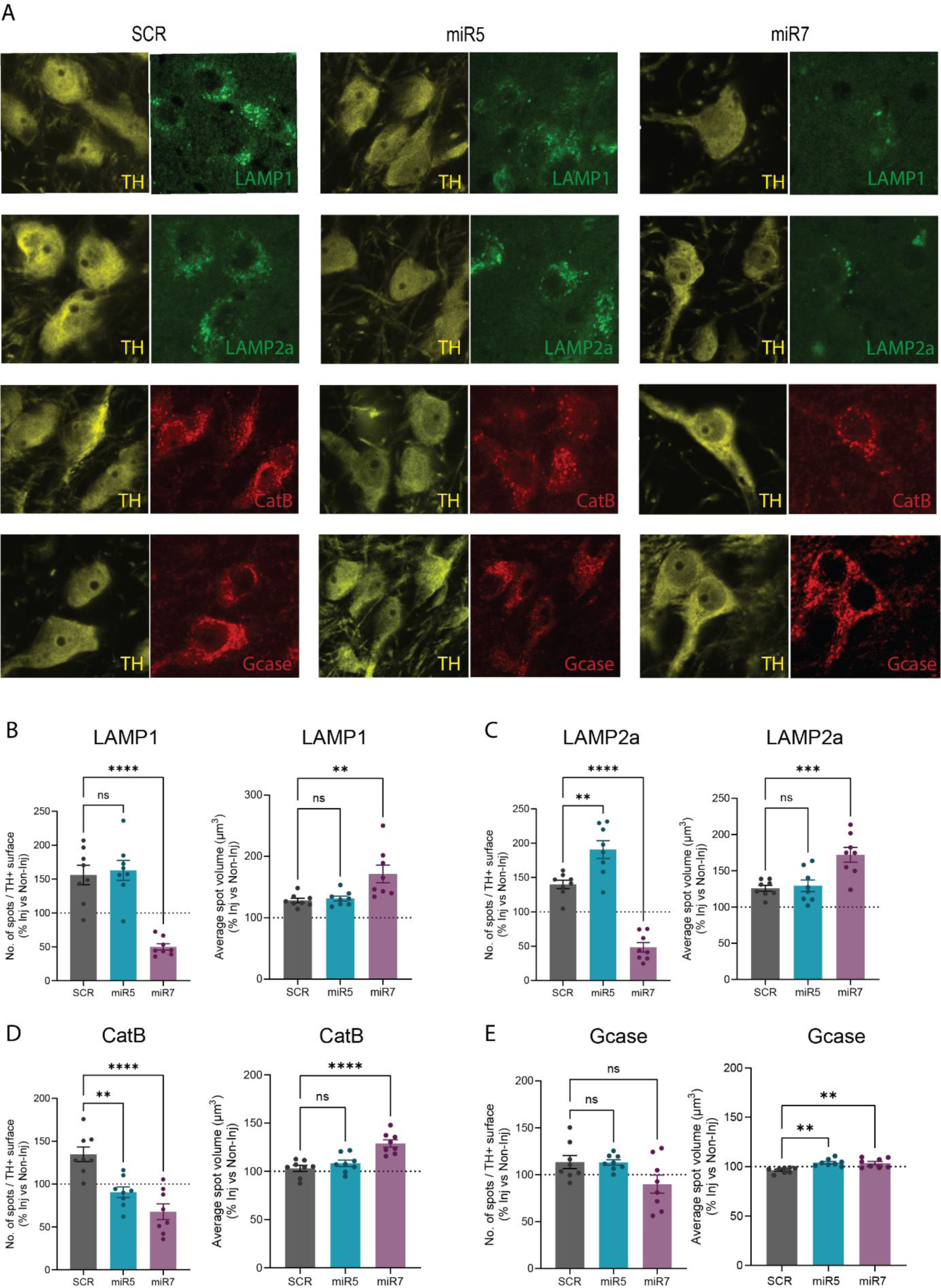
Immunofluorescence study of late endosomal/lysosomal markers in surviving dopaminergic neurons at 1 year post-injection. (A) Representative immunofluorescent images of LAMP1, LAMP2a, cathepsin B and Gcase co-stained with TH marker in the SNpc of SCR, miR5 and miR7 rats at 1 year post-injection. (B-E) Density of LAMP1+, LAMP2a+, cathepsin B+, or Gcase+ spots (number of spots per total TH+ surface) and average spot volume in TH+ cells quantified in the injected side versus non-injected side. SCR (n=8), miR5 (n=8), miR7 (n=8). (B) LAMP1+ number of spots (SCR vs. miR5 p = 0.9089, SCR vs. miR7 p < 0.0001) and LAMP1+ average spot volume (SCR vs. miR5 p > 0.9999, SCR vs. miR7 p = 0.0060). (C) LAMP2a+ number of spots (SCR vs. miR5 p = 0.0017, SCR vs. miR7 p < 0.0001) and LAMP2a+ average spot volume (SCR vs. miR5 p = 0.9435, SCR vs. miR7 p = 0.0009). (D) Cathepsin B+ number of spots (SCR vs. miR5 p = 0.0016, SCR vs. miR7 p < 0.0001) and cathepsin B+ average spot volume (SCR vs. miR5 p = 0.5022, SCR vs. miR7 p < 0.0001). (E) Gcase+ number of spots (SCR vs. miR5 p = 0.9987, SCR vs. miR7 p = 0.568) and Gcase+ average spot volume (SCR vs. miR5 p = 0.0012, SCR vs. miR7 p = 0.0020). Data are mean ± s.e.m and analyzed using one-way ANOVA followed Šídák’s multiple comparisons test, with the exception of LAMP1 average spot volume which was analyzed using Kruskal-Wallis followed Dunn’s multiple comparisons test. Each dot represents the average of 3 sections per animal.

In summary, our findings highlight the sensitivity of dopaminergic neurons to reduced ATP10B levels in rats. Knocking down ATP10B in the neurons of the SNpc leads to progressive neurodegeneration of dopaminergic neurons within this brain region, mirroring the loss of TH+ terminals in the dSTR. The surviving dopaminergic neurons in the rats injected with both ATP10B KD vectors showed a decrease in the number of cathepsin B+ organelles compared to SCR. In addition, in the group of rats injected with miR7 we could observe a reduction in the number of late endosomes/lysosomes (marked by either LAMP1, LAMP2a, or cathepsin B) together with an enlargement in the volume of these organelles in surviving dopaminergic neurons.

### ATP10B KO results in a decreased number of dopaminergic neurons in human iPSC-derived midbrain culture

To validate the susceptibility of dopaminergic neurons to the loss of ATP10B in a more translational model, ATP10B knock-out (KO) cell lines were generated in a TH-TdTomato reporter human iPSC line (BJ-SiPS) with genetic isogenic background (Fig. S5A-B). Cells were differentiated into midbrain neurons following a previously published protocol (28). On day 30 of differentiation, we evaluated the number of dopaminergic and total neurons generated. We found a significant reduction in the number of TH-positive (clone#1 p = 0.0005, clone#2 p = 0.023) (Fig. 7A, B) and MAP2-positive (p <0.0001 for clone#1-2) neurons (Fig. 7A, C) in both clones of ATP10B KO.

**Figure 7.**
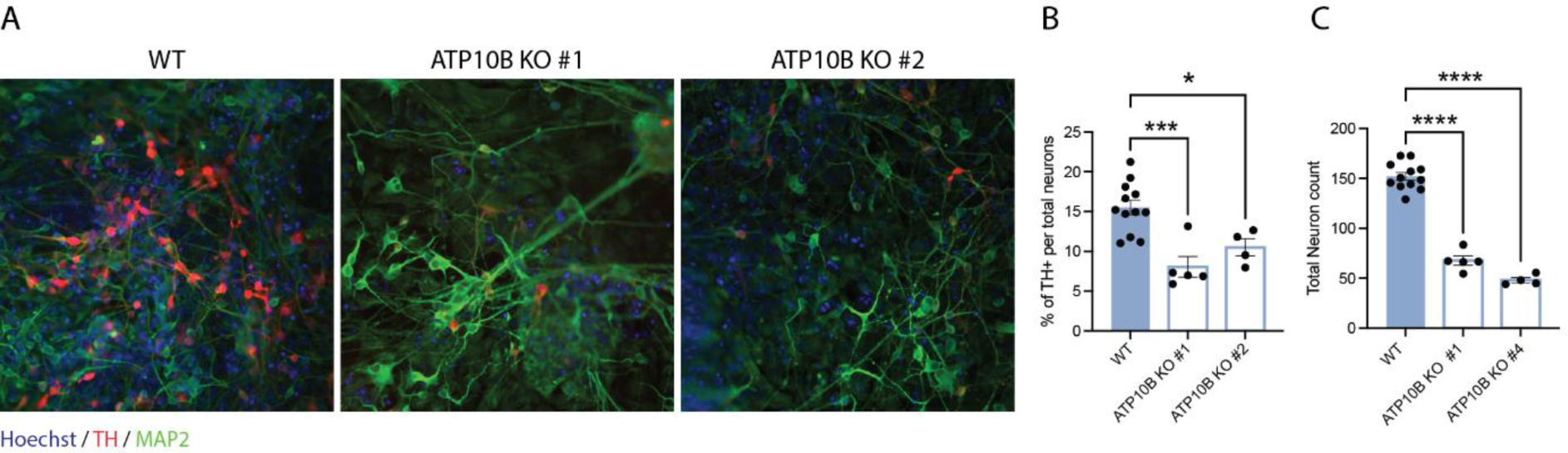
**ATP10B KO results in decreased number of dopaminergic neurons in human iPSC-derived midbrain culture**. (A) Immunocytochemistry pictures of midbrain neuronal cultures stained for TH (red) MAP2 (green) and Hoechst (blue). (B) Quantification of neuronal populations: percentage of TH+ cells over total MAP2+ population (WT vs, ATP10B KO #1 p = 0.0005, WT vs, ATP10B KO #2 p = 0.0233) and (C) total number of MAP2+ cells (WT vs, ATP10B KO #1 p < 0.0001, WT vs, ATP10B KO #2 p < 0.0001) in WT and ATP10B KO clone#1 and clone#2. Data are mean ± s.e.m and analyzed using one-way ANOVA followed by Tukey’s multiple comparison test. Each data point is the average of 3 FOV from one well.

## Discussion

ATP10B is a lysosomal transmembrane protein responsible for transporting PC and GluCer from the inner leaflet of the lysosomal membrane to the cytoplasmic side. In the context of PD, ATP10B has recently been recognized as a potential genetic risk factor, with loss of function mutations and decreased expression of ATP10B found in PD patients. Thus far, no *in vivo* model has been developed to explore the link of ATP10B to PD and SNpc dopaminergic neurons. To investigate the potential contribution of ATP10B loss of function on the nigrostriatal dopaminergic pathway, we knocked down ATP10B in the nigral neurons of adult rats.

Degeneration of the nigrostriatal pathway in individuals with PD results in progressive motor symptoms, including tremor, bradykinesia, akinesia, muscular rigidity, postural instability, and gait difficulties (1). Using rodents as animal models for PD offers advantages, as degeneration in their nigrostriatal system aligns with motor impairments that can be readily evaluated through diverse behavioral tests and experimental interventions can be more readily tested on animal models. In our experimental design, we performed unilateral injections of the knockdown AAV vector, enabling us to assess motor asymmetry deficiency as a read-out of unilateral neurodegeneration in the targeted SNpc and dysfunctional nigrostriatal dopaminergic neurotransmission. Notably, we found significant differences in preference for ipsilateral swinging in the EBST and ipsilateral rotation behavior in the open field test in both the miR5 and miR7 groups compared to the SCR group. Although only the miR5 group showed significant preferential usage of the ipsilateral forelimb in the cylinder test, the miR7 group followed a similar trend. These behaviors mirror observations in ‘hemi-parkinsonian’ models, where injections of 6-hydroxydopamine (6-OHDA) or AAV2/7 α−synuclein in the SNpc result in loss of dopaminergic neurons and asymmetric motor deficits (30–33,21). Additionally, a decrease in spontaneous locomotor activity (distance traveled and velocity) assessed in the open field test appeared to be more pronounced in the miR7 group, possibly reflecting the greater neurodegeneration of the nigrostriatal pathway observed 1 year post-injection. The miR5 rats exhibited a similar pattern, although non-significant vs. the control group. In addition, vertical exploration reflected in rearing frequency during the open field test was decreased in both the miR5 and miR7 groups. In the accelerating rotarod test for motor coordination and balance, we noted decreased performance in the ATP10B KD groups, with a stronger effect observed in the miR5 group. Interestingly, both miR5 and miR7 rats displayed muscular rigidity, as evidenced by the time spent correcting an imposed posture during the bar test. Catalepsy, commonly also associated with antipsychotic drugs, has been observed in rats bilaterally injected with 6-OHDA in the striatum, resulting in nigrostriatal dopaminergic cell loss (34). Altogether, these findings highlight behavioral deficits indicative of impaired nigrostriatal dopaminergic neurotransmission in ATP10B KD rats across different domains of motor function. There was no consistent difference between the miR5 and miR7 groups across the different behavioral tests, despite the slightly stronger dopaminergic neurodegeneration observed in the miR7 compared to the miR5 group.

In clinical practice, PET imaging of DAT using ^18^F-FE-PE2I serves as a valuable, cost-effective tool early diagnosis of neurodegenerative parkinsonism, including PD, is reimbursed as routine tool in several European countries including Belgium, and is used as an imaging marker in clinical trials (35,36). DAT PET imaging studies have revealed that PD patients show a loss of about 60% (37) of the striatal dopamine transporters by the time motor symptoms emerged, although a more recent study claimed that the loss of striatal DAT activity is about 35-45% in early PD (38). To investigate whether the loss of ATP10B in dopaminergic neurons leads to a progressive decline in striatal dopaminergic terminals, DAT PET imaging with ^18^F-FE-PE2I was employed in a sample of rats selected randomly from the larger cohort at different time points post-injection. The binding potential of the tracer to the DAT in the dopaminergic presynaptic terminals consistently showed a decrease in the ipsilateral striatum compared to the contralateral striatum in the miR5-injected rats, suggesting a progressive loss of dopaminergic terminals in the STR. Although some further DAT downregulation is observed in human neurodegenerative parkinsonism, its quantitative binding gives a reliable *in vivo* measurement of DAT availability. Because of limited access to the clinical tracer, a full longitudinal PET imaging study could not be conducted in the miR7 group. Nonetheless, we could confirm a significant decrease in the ^18^F-FE-PE2I binding in a selected group of miR7 rats at the later time points post-injection that seems to surpass the intensity of the miR5 group. Pathological examination of the brain revealed a significant loss of dopaminergic terminals (TH+) in the STR of miR5 and miR7 rats, indicating that the decrease in ^18^F-FE-PE2I binding potential occurred in parallel with a loss of striatal dopaminergic terminals.

In addition, we observed that ATP10B KD in the SNpc of rats sensitized the dopaminergic neurons to cell death at long-term following KD. The loss of both dopaminergic neurons and terminals was notably more pronounced at 1 year post-injection compared to 1 month, suggesting a progressive rather than acute impact of ATP10B loss on cellular viability, consistent with the observations from PET imaging. In order to demonstrate similar effects in a more translational model, midbrain neuronal cultures were differentiated from two different ATP10B KO human iPSC clones. In line with the results obtained in the rat model, decrease of TH positive neurons was consistently observed in both ATP10B KO clones.

Previous studies in human cell lines have linked ATP10B KD to a loss of LAMP1+ lysosomal mass, decreased lysosomal pH, reduced lysosomal degradative capacity, and impaired lysosomal membrane integrity upon rotenone exposure. In primary neurons, ATP10B KD not only decreased lysosomal degradative capacity but also increased susceptibility to cell death (9). Lysosomal activity and autophagy pathways are essential in maintaining neuronal health and proper α−synuclein levels in neurons (39). α−synuclein aggregation and accumulation in Lewy bodies, together with a variety of membrane components such as abnormal lysosomes or mitochondria, is one of the main hallmarks of PD (6). Although we were unable to detect a difference in total α−synuclein or lysosomal protein levels measured by western blot in tissue extracts of the SN, immunofluorescence staining revealed changes in late endosomal/lysosomal markers in the surviving dopaminergic neurons of rats injected with ATP10B KD vectors at 1 year post-injection. Particularly there was a reduction in the number of late endosomes/lysosomes and the enlargement of these organelles within surviving dopaminergic neurons of rats injected with miR7. However, the pattern observed in the miR5 group of rats was different, so further experiments are needed to explore the link between ATP10B loss and lysosomal pathways alteration in dopaminergic neurons. It is know that the soma of neurons is the primary site for cargo degradation. Degradative lysosomes are observed to be more abundant in the soma of neurons compared to dendrites. Additionally, degradative autolysosomes, originating from distal axonal regions, retrogradely travel to the soma (40). However, lysosomes also participate in essential processes in presynaptic terminals, such as autophagy and neurotransmitter recycling (41–45). Dopaminergic neurons display extensive and complex axonal arborization, with an increased need of ATP and increased production of reactive oxygen species (ROS), which make them vulnerable to neurodegeneration (46). Although not specifically evaluated, this information may suggest that impairment of lysosomal activity caused by ATP10B loss of function could potentially lead to impairments not only in the soma but also at the level of the presynaptic terminals, affecting the maintenance of the nigrostriatal dopaminergic pathway with age.

Biochemical experiments have shown that disease-associated ATP10B variants, as well as KD of the protein, result in the loss of the ability to translocate PC and GluCer (9,17). Interestingly, ATP10B shares the same substrate with the enzyme Gcase, responsible for breaking down GluCer into ceramide and glucose within lysosomes. Gcase is encoded by GBA, the main genetic risk factor in PD (18). The metabolism of ceramides and GluCer seems to play a crucial role in PD, as evidenced by elevated levels in the plasma of sporadic PD patients, which correlate with the severity of cognitive impairment (47,48). *In vitro* studies conducted on iPSC-derived dopaminergic neurons have demonstrated that decreased Gcase protein levels and activity lead to elevated GluCer levels, compromising lysosomal function and autophagy, and triggering a senescence-like phenotype in these neurons (49,50). In addition, not only GluCer but accumulation of other lipids has been reported in dopaminergic neurons of PD patients (51,52). Given these findings, investigating GluCer/lipid levels in dopaminergic neurons following ATP10B loss could provide valuable insights into potential analogous alterations in our *in vivo* model.

Previously, other proteins from the P-type ATPases have been identified in a broad range of diseases (53), including loss of functions mutations in ATP13A2 as cause for the severe parkinsonism Kufor-Rakeb syndrome (KRS) (54,55). ATP10B has emerged as a new potential genetic risk factor in PD and previous *in vitro* studies have provided insight into the function of ATP10B and the repercussions of its loss on the lysosomes and at cellular level (9,17). Our study establishes for the first time a link between ATP10B and PD in an *in vivo* model. Our findings demonstrate that the depletion of ATP10B in the neurons of the SNpc leads to progressive neurodegeneration of dopaminergic neurons and the loss of dopaminergic terminals in the STR, accompanied by parkinsonian motor impairments in rats. In forthcoming experiments, it will be interesting to explore in detail whether lysosomal impairments and/or accumulation of ATP10B substrates may underlie the observed nigrostriatal pathology.

## ACKNOWLEDGEMENTS AND FUNDING

We thank Annelies Aertgeerts, Krist’l Vennekens and Joris Van Asselberghs for their technical assistance and the Leuven Viral Vector Core (LVVC) for the rAAV vector production. We acknowledge the VIB Bioimaging Core for the use of the confocal microscope and technical assistance. This research was funded by the Aligning Science Across Parkinson’s [ASAP-000458] through the Michael J. Fox Foundation for Parkinson’s Research (MJFF) allocated to PV and VB, the Vlaio O&O project LeAD-UP, and the KU Leuven Parkinson fund. EB is a postdoctoral research fellow supported by Aligning Science Across Parkinson’s [ASAP-000458]. TT-M, DC and GT were supported by doctoral fellowships from the Research Foundation-Flanders (FWO). CC was funded by FWO I000321N and FWO I000123N, and the PET-CT equipment was funded via an FWO medium infrastructure grant (AKUL15-30/G0H1216N) allocated to KVL. KVL was funded by a senior clinical researcher mandate of FWO. The authors would like to thank the staff of the clinical radiopharmacy at UZ Leuven for supply of [^18^F]-FE-PE2I, and Ms. Kasia Błażejczyk for technical support during some of the *in vivo* imaging assays.

## AUTHORS’ CONTRIBUTIONS

MSC, EB, VB, PV, and JB conceived the project. MSC, EB, VB, EC, CC, GMP, CVH, and JB designed the experiments. MSC, EB, EC, CC, GMP, TTM, DC, and GT conducted the experiments. MSC, EB, VB, EC, CC, GMP, KVL, TTM, AC, CVH, PV, and JB analyzed data and interpreted results. MSC, EB, VB, EC, CC, and GMP wrote the paper. All authors read and approved the final manuscript.

## DATA AVAILABILITY

All datasets generated and analyzed during this study are available in the Zenodo repository (10.5281/zenodo.12699264).

## ETHICAL APPROVAL AND CONSENT TO PARTICIPATE

Not applicable

## CONSENT FOR PUBLICATION

Not applicable

## ABBREVIATIONS

PD: Parkinson’s disease
SNpc: substantia nigra pars compacta
dSTR: dorsal striatum
DLB: dementia with Lewy bodies
GluCer: glucosylceramide
PC: phosphatidylcholine
KD: knockdown
Gcase: glucocerebrosidase
AAV: adeno-associated vector
DAT: dopamine transporter
PET: positron emission tomography imaging
EBST: elevated body swing test
CT: computered tomography
MLEM: maximum-likelihood expectation-maximization
Hus: hounsfield units
SRTM: standard reference tissue model
BPnd: binding potential
PFA: paraformaldehyde
PBS-T: PBS with 0.1% Tergitol
TH: tyrosine hydroxylase
PVDF: polyvinylidene fluoride
hiPSCs: Human induced pluripotent stem cells
KO: knock-out
6-OHDA: 6-hydroxydopamine
ROS: reactive oxygen species
KRS: Kufor-Rakeb syndrome

---

**Figure S1.**
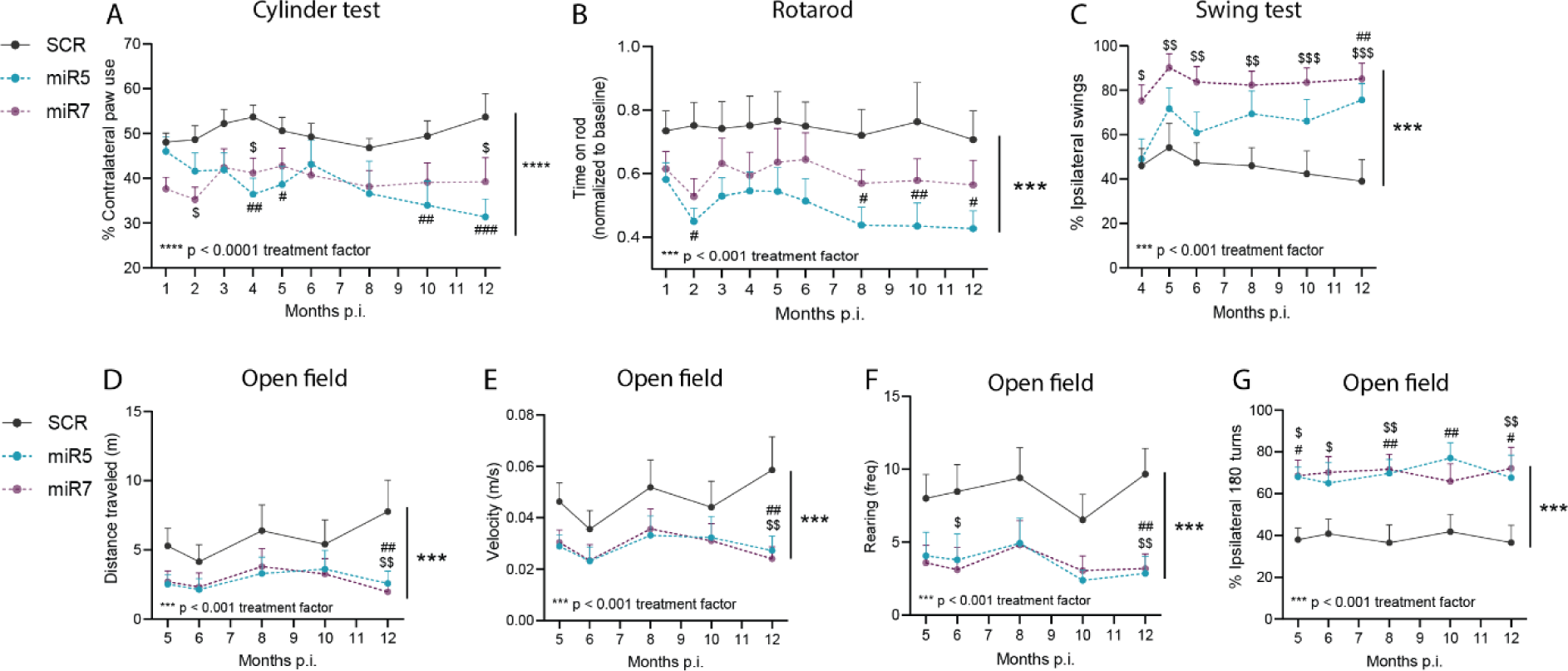
ATP10B KD in SNpc neurons leads to motor asymmetry and decreased motor function. (A, C, G) Indicative of decreased ipsilateral dopaminergic neurotransmission, rats with ATP10B KD showed decreased preference in the use of the contralateral paw in the cylinder test (A), increased bias to ipsilateral swing in the elevated body swing test (C), and spontaneous ipsilateral turning in the open field test (G). (B) Motor coordination and balance was assessed using an accelerated rotarod test (4-40 rpm), with the average performance post-lesion normalized to the individual rat performance prior to surgery. (D, E, F) Spontaneous motor behavior was recorded during a 5-min open field test and analyzed for total distance traveled (D), velocity (E) and rearing (F). Data are mean + s.e.m. *** p < 0.001 (two-way ANOVA, treatment factor), # p < 0.05, ## p < 0.01, ### p < 0.001 (Dunnett’s post hoc test miR5 vs. SCR at corresponding time point), $ p < 0.05, $$ p < 0.01, $$$ p < 0.001 (Dunnett’s post hoc test miR7 vs. SCR at corresponding time point).

**Figure S2.**
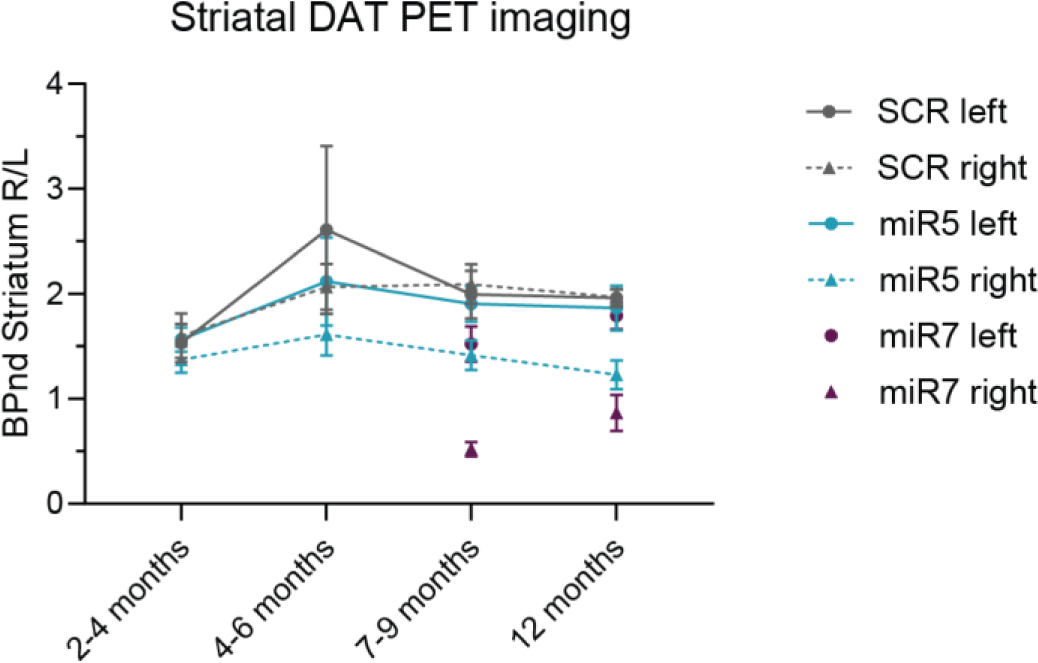
Raw BPnd values from 18F-FE-PE2I PET imaging. Striatal DAT binding potential (BPnd), measured by 18F-FE-PE2I microPET imaging, in the ipsilateral (right) and the contralateral (left) striatum observed in miR5-injected animals (n=5), SCR-injected animals (n=3) and miR7-injected animals at 7-9 months (n=2) and 12 months (n=3).

**Figure S3.**
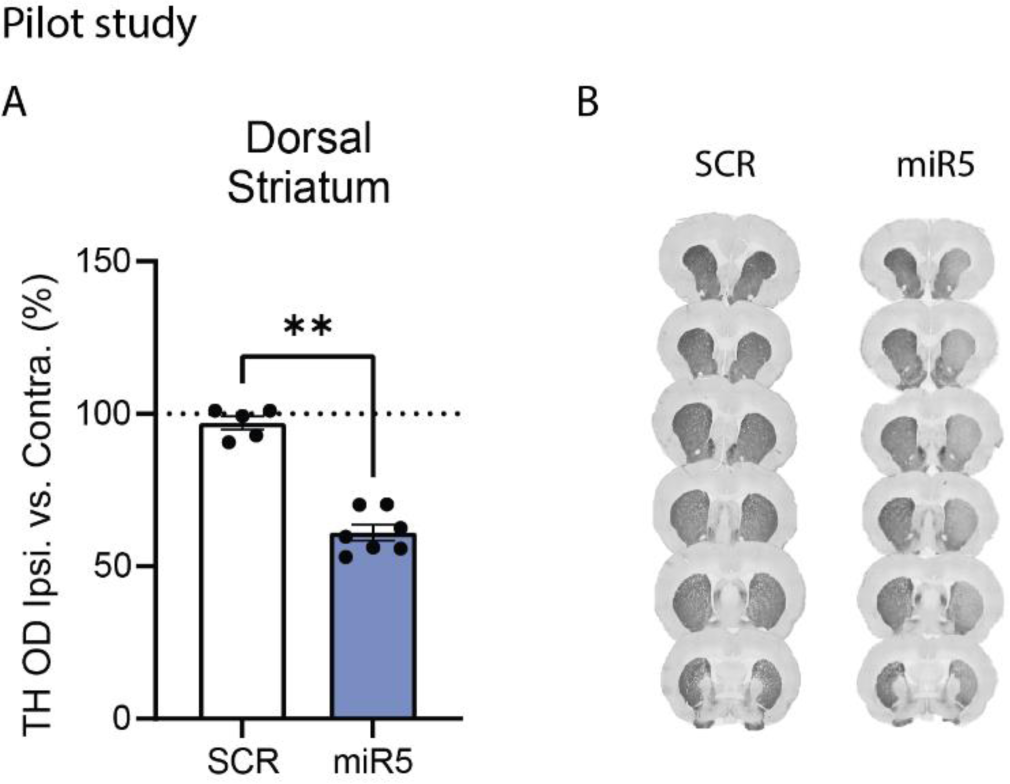
Pilot study: ATP10B KD leads to decreased dopaminergic terminals in the dSTR of miR5 group. TH positive area was quantified using ImageJ in 6 different sections that cover the STR. (A) Percentage of TH positive area ipsilateral (R) versus contralateral (L) dorsal STR of 1 year post-injection rats, SCR (n=5) and miR5 (n=7). Data are mean ± s.e.m and analyzed using t test, ** p = 0.0025. Each dot represents the average of the 6 sections per animal. (B) Representative image of TH immunohistochemical staining on 6 different sections from one SCR and miR5 rat 1 year post-injection.

**Figure S4.**
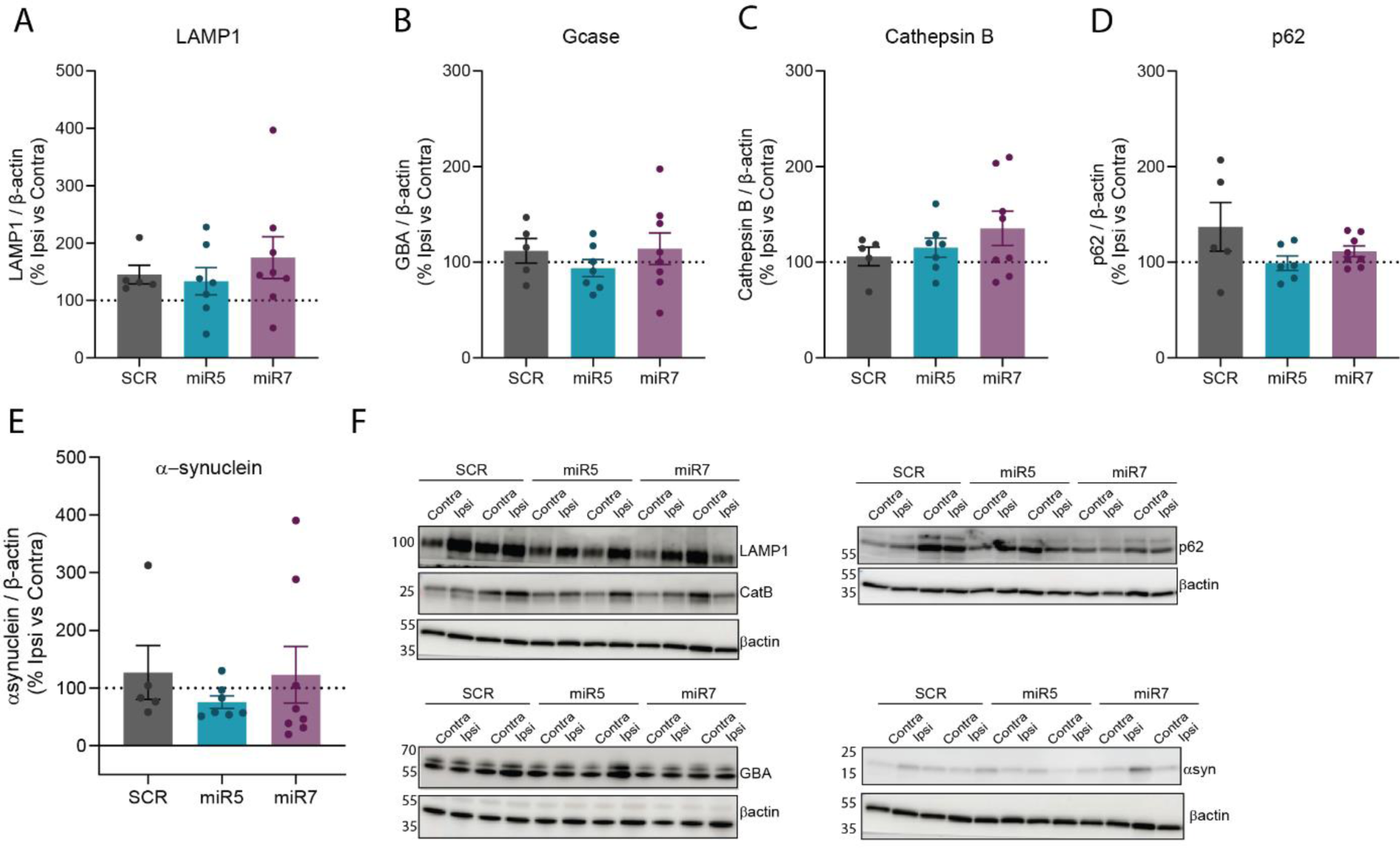
No significant changes in lysosomal proteins or total *α*-synuclein levels in whole SN protein extracts. (A-E) Protein signal values were normalized to the endogenous protein (β-actin) levels, and the ipsilateral (Ipsi) versus contralateral (Contra) percentage of the protein signal is represented. Data are presented as mean ± s.e.m., with each dot representing an individual animal. (F) Representative western blot images of α-synuclein and lysosomal proteins analyzed in whole SN extracts from SCR, miR5, and miR7 rats.

**Figure S5.**
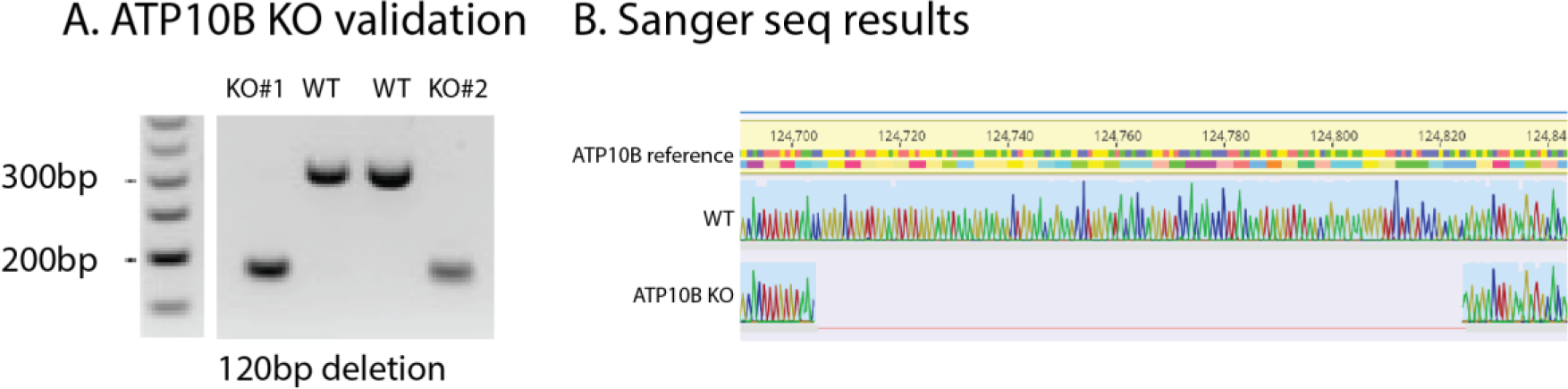
ATP10B knock-out (KO) cell lines generated in a TH-TdTomato reporter hiPSC line (BJ-SiPS). (A) Electrophoresis gel image after PCR showing the loss of a 120 bp fragment after CRISPR KO targeting exon 1 of ATP10B. (B) Sanger sequencing results confirming the deletion of the same 120 bp fragment.

## References

1. Bloem BR, Okun MS, Klein C. Parkinson’s disease. The Lancet. 2021 Jun;397(10291):2284–303.

2. Kamath T, Abdulraouf A, Burris SJ, Langlieb J, Gazestani V, Nadaf NM, et al. Single-cell genomic profiling of human dopamine neurons identifies a population that selectively degenerates in Parkinson’s disease. Nat Neurosci. 2022 May;25(5):588–95.

3. Andén NE, Carlsson A, Dahlström A, Fuxe K, Hillarp NÅ, Larsson K. Demonstration and mapping out of nigro-neostriatal dopamine neurons. Life Sci. 1964 Jun 1;3(6):523–30.

4. Redgrave P, Rodriguez M, Smith Y, Rodriguez-Oroz MC, Lehericy S, Bergman H, et al. Goal-directed and habitual control in the basal ganglia: implications for Parkinson’s disease. Nat Rev Neurosci. 2010 Nov;11(11):760–72.

5. Cramb KML, Beccano-Kelly D, Cragg SJ, Wade-Martins R. Impaired dopamine release in Parkinson’s disease. Brain. 2023 Aug 1;146(8):3117–32.

6. Shahmoradian SH, Lewis AJ, Genoud C, Hench J, Moors TE, Navarro PP, et al. Lewy pathology in Parkinson’s disease consists of crowded organelles and lipid membranes. Nat Neurosci. 2019 Jul;22(7):1099–109.

7. Navarro-Romero A, Montpeyó M, Martinez-Vicente M. The Emerging Role of the Lysosome in Parkinson’s Disease. Cells. 2020 Nov 2;9(11):2399.

8. Malpartida AB, Williamson M, Narendra DP, Wade-Martins R, Ryan BJ. Mitochondrial Dysfunction and Mitophagy in Parkinson’s Disease: From Mechanism to Therapy. Trends Biochem Sci. 2021 Apr;46(4):329–43.

9. Martin S, Smolders S, Van den Haute C, Heeman B, van Veen S, Crosiers D, et al. Mutated ATP10B increases Parkinson’s disease risk by compromising lysosomal glucosylceramide export. Acta Neuropathol (Berl). 2020 Jun 1;139(6):1001–24.

10. Zhao Y, Pan H, Wang Y, Zeng Q, Fang Z, He R, et al. ATP10B variants in Parkinson’s disease: a large cohort study in Chinese mainland population. Acta Neuropathol (Berl). 2021 May 1;141(5):805–6.

11. Tesson C, Lohmann E, Devos D, Bertrand H, Lesage S, Brice A. Segregation of ATP10B variants in families with autosomal recessive parkinsonism. Acta Neuropathol (Berl). 2020 Nov;140(5):783–5.

12. Real R, Moore A, Blauwendraat C, Morris HR, Bandres-Ciga S, the International Parkinson’s Disease Genomics Consortium (IPDGC). ATP10B and the risk for Parkinson’s disease. Acta Neuropathol (Berl). 2020 Sep 1;140(3):401–2.

13. Díaz-Belloso R, Muñoz-Delgado L, Martín-Bornez M, Ojeda E, Periñán MT, García-Díaz S, et al. Role of ATP10B in Parkinson disease in a cohort from southern Spain. Parkinsonism Relat Disord. 2024 Jul;124:106989.

14. Smolders S, Van Broeckhoven C. Reply: Segregation of ATP10B variants in families with autosomal recessive Parkinsonism. Acta Neuropathol (Berl). 2020 Nov;140(5):787–9.

15. Smolders S, Van Broeckhoven C. Reply: ATP10B variants in Parkinson’s disease—a large cohort study in Chinese mainland population. Acta Neuropathol (Berl). 2021;141(5):807–8.

16. Smolders S, Van Broeckhoven C. Reply: ATP10B and the risk for Parkinson’s disease. Acta Neuropathol (Berl). 2020 Sep 1;140(3):403–4.

17. Wouters R, Beletchi I, Van den Haute C, Baekelandt V, Martin S, Eggermont J, et al. The lipid flippase ATP10B enables cellular lipid uptake under stress conditions. Biochim Biophys Acta BBA - Mol Cell Res. 2024 Feb 1;1871(2):119652.

18. Sidransky E, Lopez G. The link between the GBA gene and parkinsonism. Lancet Neurol. 2012 Nov 1;11(11):986–98.

19. van Meer G, Voelker DR, Feigenson GW. Membrane lipids: where they are and how they behave. Nat Rev Mol Cell Biol. 2008 Feb;9(2):112–24.

20. Clarke RJ, Hossain KR, Cao K. Physiological roles of transverse lipid asymmetry of animal membranes. Biochim Biophys Acta BBA - Biomembr. 2020 Oct 1;1862(10):183382.

21. Van der Perren A, Toelen J, Casteels C, Macchi F, Van Rompuy AS, Sarre S, et al. Longitudinal follow-up and characterization of a robust rat model for Parkinson’s disease based on overexpression of alpha-synuclein with adeno-associated viral vectors. Neurobiol Aging. 2015 Mar 1;36(3):1543–58.

22. Gulyás M, Bencsik N, Pusztai S, Liliom H, Schlett K. AnimalTracker: An ImageJ-Based Tracking API to Create a Customized Behaviour Analyser Program. Neuroinformatics. 2016 Oct;14(4):479–81.

23. In vitro autoradiography and in vivo evaluation in cynomolgus monkey of [18F]FE-PE2I, a new dopamine transporter PET radioligand - PubMed [Internet]. [cited 2024 Jul 8]. Available from: https://pubmed.ncbi.nlm.nih.gov/19562698/

24. Iterative CT Reconstruction Using Shearlet-Based Regularization | IEEE Journals & Magazine | IEEE Xplore [Internet]. [cited 2024 Feb 27]. Available from: https://ieeexplore.ieee.org/document/6589007

25. Wk S, Mm M, A B, Dl A, V P, Sl D. Serial microPET measures of the metabolic reaction to a microdialysis probe implant. J Neurosci Methods [Internet]. 2006 Sep 15 [cited 2024 Jul 8];155(2). Available from: https://pubmed.ncbi.nlm.nih.gov/16519945/

26. Yamasaki T, Fujinaga M, Kawamura K, Furutsuka K, Nengaki N, Shimoda Y, et al. Dynamic Changes in Striatal mGluR1 But Not mGluR5 during Pathological Progression of Parkinson’s Disease in Human Alpha-Synuclein A53T Transgenic Rats: A Multi-PET Imaging Study. J Neurosci Off J Soc Neurosci. 2016 Jan 13;36(2):375–84.

27. Rosas-Arellano A, Villalobos-González JB, Palma-Tirado L, Beltrán FA, Cárabez-Trejo A, Missirlis F, et al. A simple solution for antibody signal enhancement in immunofluorescence and triple immunogold assays. Histochem Cell Biol. 2016 Oct 1;146(4):421–30.

28. Kim TW, Piao J, Koo SY, Kriks S, Chung SY, Betel D, et al. Biphasic Activation of WNT Signaling Facilitates the Derivation of Midbrain Dopamine Neurons from hESCs for Translational Use. Cell Stem Cell. 2021 Feb 4;28(2):343–355.e5.

29. Taylor TN, Greene JG, Miller GW. Behavioral phenotyping of mouse models of Parkinson’s disease. Behav Brain Res. 2010 Jul 29;211(1):1–10.

30. Hefti F, Melamed E, Wurtman RJ. Partial lesions of the dopaminergic nigrostriatal system in rat brain: biochemical characterization. Brain Res. 1980 Aug 11;195(1):123–37.

31. Perese DA, Ulman J, Viola J, Ewing SE, Bankiewicz KS. A 6-hydroxydopamine-induced selective parkinsonian rat model. Brain Res. 1989 Aug 14;494(2):285–93.

32. Schwarting RK, Huston JP. The unilateral 6-hydroxydopamine lesion model in behavioral brain research. Analysis of functional deficits, recovery and treatments. Prog Neurobiol. 1996 Oct;50(2–3):275–331.

33. Schober A. Classic toxin-induced animal models of Parkinson’s disease: 6-OHDA and MPTP. Cell Tissue Res. 2004 Oct 1;318(1):215–24.

34. Lindner MD, Cain CK, Plone MA, Frydel BR, Blaney TJ, Emerich DF, et al. Incomplete nigrostriatal dopaminergic cell loss and partial reductions in striatal dopamine produce akinesia, rigidity, tremor and cognitive deficits in middle-aged rats. Behav Brain Res. 1999 Jul 1;102(1):1– 16.

35. Schou M, Steiger C, Varrone A, Guilloteau D, Halldin C. Synthesis, radiolabeling and preliminary in vivo evaluation of [18F]FE-PE2I, a new probe for the dopamine transporter. Bioorg Med Chem Lett. 2009 Aug 15;19(16):4843–5.

36. Kerstens VS, Fazio P, Sundgren M, Brumberg J, Halldin C, Svenningsson P, et al. Longitudinal DAT changes measured with [18F]FE-PE2I PET in patients with Parkinson’s disease; a validation study. NeuroImage Clin. 2023 Feb 11;37:103347.

37. Ribeiro MJ, Vidailhet M, Loc’h C, Dupel C, Nguyen JP, Ponchant M, et al. Dopaminergic function and dopamine transporter binding assessed with positron emission tomography in Parkinson disease. Arch Neurol. 2002 Apr;59(4):580–6.

38. Heng N, Malek N, Lawton MA, Nodehi A, Pitz V, Grosset KA, et al. Striatal Dopamine Loss in Early Parkinson’s Disease: Systematic Review and Novel Analysis of Dopamine Transporter Imaging. Mov Disord Clin Pract. 2023 Feb 17;10(4):539–46.

39. Vogiatzi T, Xilouri M, Vekrellis K, Stefanis L. Wild type alpha-synuclein is degraded by chaperone-mediated autophagy and macroautophagy in neuronal cells. J Biol Chem. 2008 Aug 29;283(35):23542–56.

40. Kulkarni VV, Maday S. Neuronal endosomes to lysosomes: A journey to the soma. J Cell Biol. 2018 Sep 3;217(9):2977–9.

41. Sambri I, D’Alessio R, Ezhova Y, Giuliano T, Sorrentino NC, Cacace V, et al. Lysosomal dysfunction disrupts presynaptic maintenance and restoration of presynaptic function prevents neurodegeneration in lysosomal storage diseases. EMBO Mol Med. 2017 Jan;9(1):112–32.

42. Soukup S, Vanhauwaert R, Verstreken P. Parkinson’s disease: convergence on synaptic homeostasis. EMBO J. 2018 Sep 14;37(18):e98960.

43. Nguyen M, Wong YC, Ysselstein D, Severino A, Krainc D. Synaptic, Mitochondrial, and Lysosomal Dysfunction in Parkinson’s Disease. Trends Neurosci. 2019 Feb;42(2):140–9.

44. Cunha Sr. CAV, Burrinha T, Almeida CG. Lysosomal subcellular distribution in neurons: Are there synaptic lysosomes? Alzheimers Dement. 2021;17(S3):e053844.

45. Sulzer D, Bogulavsky J, Larsen KE, Behr G, Karatekin E, Kleinman MH, et al. Neuromelanin biosynthesis is driven by excess cytosolic catecholamines not accumulated by synaptic vesicles. Proc Natl Acad Sci U S A. 2000 Oct 24;97(22):11869–74.

46. Pacelli C, Giguère N, Bourque MJ, Lévesque M, Slack RS, Trudeau LÉ. Elevated Mitochondrial Bioenergetics and Axonal Arborization Size Are Key Contributors to the Vulnerability of Dopamine Neurons. Curr Biol. 2015 Sep 21;25(18):2349–60.

47. Mielke MM, Maetzler W, Haughey NJ, Bandaru VVR, Savica R, Deuschle C, et al. Plasma Ceramide and Glucosylceramide Metabolism Is Altered in Sporadic Parkinson’s Disease and Associated with Cognitive Impairment: A Pilot Study. Blum D, editor. PLoS ONE. 2013 Sep 18;8(9):e73094.

48. Huh YE, Park H, Chiang MSR, Tuncali I, Liu G, Locascio JJ, et al. Glucosylceramide in cerebrospinal fluid of patients with GBA-associated and idiopathic Parkinson’s disease enrolled in PPMI. Npj Park Dis. 2021 Nov 22;7(1):1–7.

49. Schöndorf DC, Aureli M, McAllister FE, Hindley CJ, Mayer F, Schmid B, et al. iPSC-derived neurons from GBA1-associated Parkinson’s disease patients show autophagic defects and impaired calcium homeostasis. Nat Commun. 2014 Jun 6;5(1):4028.

50. Russo T, Kolisnyk B, B. S. A, Plessis-Belair J, Kim TW, Martin J, et al. The SATB1-MIR22-GBA axis mediates glucocerebroside accumulation inducing a cellular senescence-like phenotype in dopaminergic neurons. Aging Cell. n/a(n/a):e14077.

51. Plotegher N, Bubacco L, Greggio E, Civiero L. Ceramides in Parkinson’s Disease: From Recent Evidence to New Hypotheses. Front Neurosci [Internet]. 2019 [cited 2024 Feb 26];13. Available from: https://www.frontiersin.org/journals/neuroscience/articles/10.3389/fnins.2019.00330

52. Brekk OR, Honey JR, Lee S, Hallett PJ, Isacson O. Cell type-specific lipid storage changes in Parkinson’s disease patient brains are recapitulated by experimental glycolipid disturbance. Proc Natl Acad Sci. 2020 Nov 3;117(44):27646–54.

53. Van der Mark VA, Elferink RPJO, Paulusma CC. P4 ATPases: Flippases in Health and Disease. Int J Mol Sci. 2013 Apr;14(4):7897–922.

54. Di Fonzo A, Chien HF, Socal M, Giraudo S, Tassorelli C, Iliceto G, et al. ATP13A2 missense mutations in juvenile parkinsonism and young onset Parkinson disease. Neurology. 2007 May 8;68(19):1557–62.

55. Ramirez A, Heimbach A, Gründemann J, Stiller B, Hampshire D, Cid LP, et al. Hereditary parkinsonism with dementia is caused by mutations in ATP13A2, encoding a lysosomal type 5 P-type ATPase. Nat Genet. 2006 Oct;38(10):1184–91.

